# Increasing transposase abundance with ocean depth correlates with a particle-associated lifestyle

**DOI:** 10.1101/2022.04.27.489810

**Authors:** Juntao Zhong, Thais del Rosario Hernández, Oleksandr Kyrysyuk, Benjamin J. Tully, Rika E. Anderson

## Abstract

Transposases are mobile genetic elements (MGEs) that move within and between genomes, promoting genomic plasticity in microorganisms. In marine microbial communities, the abundance of transposases increases with depth, but the reasons behind this trend remain unclear. Our analysis of metagenomes from the *Tara* Oceans and Malaspina Expeditions suggests that a particle-associated lifestyle is the main covariate for the high occurrence of transposases in the deep ocean. We observed a strong and depth-independent correlation between transposase abundance and the presence of biofilm-associated genes, as well as the prevalence of secretory enzymes. This suggests that mobile genetic elements readily propagate among microbial communities within crowded biofilms. Furthermore, we show that particle association positively correlates with larger genome size, which is in turn is associated with higher transposase abundance. Cassette sequences associated with transposons are enriched with genes related to defense mechanisms, which are more highly expressed in the deep sea. Thus, while transposons spread at the expense of their microbial hosts, they also introduce novel genes and potentially benefit the hosts in helping to compete for limited resources. Overall, our results suggest a new understanding of deep ocean particles as highways for gene sharing among defensively oriented microbial genomes.

**Importance:** Genes can move within and between microbial genomes via mobile genetic elements, which include transposases and transposons. In the oceans, there is a puzzling increase in transposase abundance in microbial genomes as depth increases. To gain insight into this trend, we conducted an extensive analysis of marine microbial metagenomes and metatranscriptomes. We found a significant correlation between transposase abundance and a particle-associated lifestyle among marine microbes. We also observed a link between transposase abundance and genes related to defense mechanisms. These results suggest that as microbes become densely packed into crowded particles, mobile genes are more likely to spread and carry genetic material that provides a competitive advantage in crowded habitats. This may enable deep sea microbes to effectively compete in such environments.

## Introduction

Mobile genetic elements (MGEs) are segments of DNA that facilitate the movement of genetic sequences within and between bacterial and archaeal genomes (1). MGEs generally encode enzymes to mediate this process (1): in transposons, the mediating enzymes are transposases, and the mechanism of transfer results in inverted repeats (2, 3). During migration, some MGEs carry cassette sequences that often contain functional genes, including those for antibiotic resistance (4), metal uptake (5, 6), and regulatory genes influencing gene expression in host cells (7, 8).

Transposases are the most abundant and ubiquitous genes in nature (9), although their distribution among taxa is uneven (10, 11). In marine systems, one of the most striking – and as of yet unexplained – trends in microbial genomics is the distinct increase in transposase abundance with depth. Genomic analyses of samples collected at station ALOHA in the North Pacific revealed a substantial increase in transposase abundance from 500 m to their observed maximum at 4000 m (12, 13). Transposases were one of the most overrepresented COG-categories in ALOHA deep waters, accounting for 1.2% of all fosmid sequences from 4000 m (12, 14). Similarly, a study of hydrothermal chimneys demonstrated a high abundance of transposases in biofilms, comprising 8% of all metagenomic reads, which is ten times higher than observed in metagenomes from in other habitats (15). Based on these observations, our analyses focused on three central questions: 1) why does transposase abundance increase with depth in marine systems, 2) are transposases selfish genes with neutral or deleterious effects on host genomes, and 3) do transposases provide useful functions to the microbial hosts that harbor them?

To better understand the high transposase abundance in the deep sea and to gain insights into the role of MGEs in marine microbial communities, we analyzed 138 microbial metagenomes and 152 microbial metatranscriptomes from the *Tara* Oceans Expedition (16, 17). The *Tara* Oceans samples spanned depths from 5m to 1000 m. In order to represent bathypelagic microbial communities, we also incorporated 58 metagenomes from the 2010 Malaspina Expedition (18), collected between 2400m and 4000m. To explore the role of transposases at a genome-resolved level, we analyzed 1402 metagenome assembled genomes (MAGs) previously generated from the *Tara* Oceans (19) and Malaspina (18) metagenomes. Previous studies have shown that deep-sea prokaryotes have a predominantly particle-associated lifestyle (20). Here, we show that the increasing abundance of transposases with depth in ocean microbial communities is associated with a shift towards a particle-associated lifestyle. We identified taxonomy, particle association, and genome size as key covariates with transposase abundance in MAGs. Additionally, we observed a high abundance of ORFs in the functional category “defense mechanisms” among the open reading frames (ORFs) in cassette sequences associated with transposases, suggesting that transposons introduce beneficial genes to their microbial hosts inhabiting highly competitive habitats.

## Results and Discussion

Previous studies have revealed that transposase abundance in microbial genomes is associated with increased ocean depth, lower dissolved oxygen (DO) concentrations, and a particle-associated lifestyle (as opposed to a planktonic lifestyle) (10, 13, 21). However, the reason and nature of these correlations has not been further explored. To confirm these findings on a broader scale, we screened for significant covariates for transposase abundance in the *Tara* Oceans and Malaspina metagenomes. We plotted the transposase abundance against depth and DO. In our metagenome analysis, transposase abundance was defined as the proportion of reads mapped back to all transposase ORFs (the database used to identify putative transposase ORFs is shown in **Table S1**). We confirmed that transposase abundance increases steadily as depth increases, despite large variations across samples (**Fig. 1**). In every ocean, the transposase abundances of the deep water samples were higher than those of the shallow water samples (**Table S2**). Here, “deep water” was defined as mesopelagic (depth 250-1000 m) and bathypelagic (2500-4000 m) waters, and “shallow water” was defined as surface (<10 m) and deep chlorophyll maximum (DCM, depth 17-120 m) waters.

**Fig. 1:**
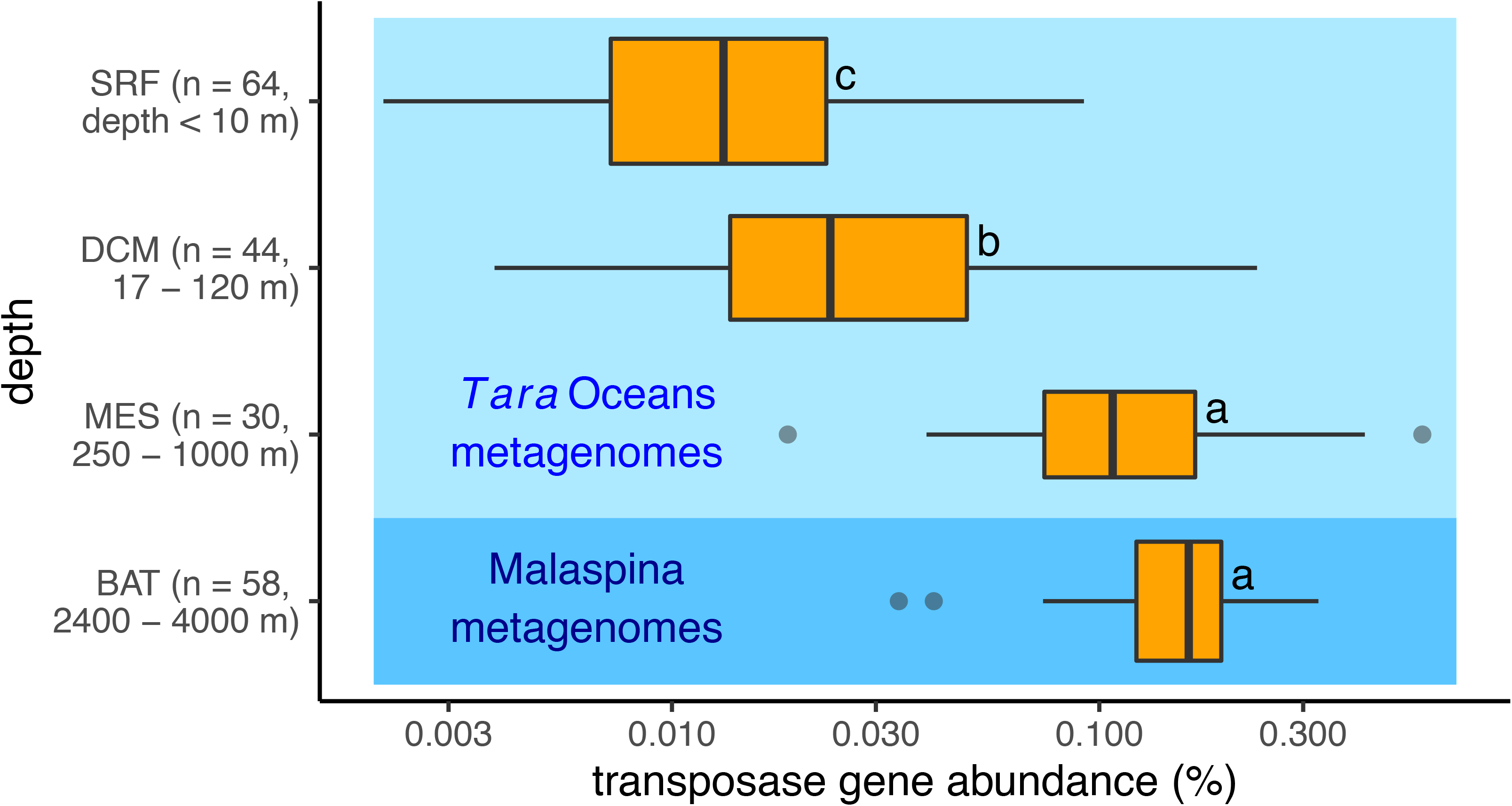
Samples from the deep ocean have higher transposase abundance. The abundance of transposase ORFs in metagenomic samples from the *Tara* Oceans and Malaspina Expedition, separated by depth. SRF: surface; DCM: deep chlorophyll maximum; MES: mesopelagic zone; BAT: bathypelagic zone. (a), (b) and (c) represent Tukey’s honestly significant differences, with each letter representing groups that are statistically significant.

The depth of a sample (e.g., surface, DCM) was a significant predictor for transposase abundance (Table S3). Adding temperature, however, did not significantly improve the prediction of transposase abundance in a sample (ANOVA *F*-test *p* =0.48, **Table S3** and **Fig. S1A**). Similarly, adding dissolved oxygen also did not improve the prediction of transposase abundance (ANOVA *F*-test *p* = 0.09, **Table S3**). This was partially due to the inconsistent relationship between DO and transposase abundance across depths. In the surface and bathypelagic samples, DO positively correlated with transposase abundance, but this correlation was reversed in DCM and mesopelagic samples (**Fig. S1B**).

### Transposase abundance is positively correlated with a particle-associated lifestyle in all depths

To further investigate the relationship between transposase abundance and particle-associated lifestyles, we examined the correlation between transposase abundance and the percentage of predicted secretory carbohydrate-active enzymes (CAZymes) and peptidases among all CAZyme and peptidase ORFs (see Methods). The percentage of secretory enzymes among all CAZymes and peptidases has previously been applied to quantify the degree to which microbial communities rely on a particle-associated lifestyle (20). Microorganisms rely on extracellular enzymes to degrade large particulate organic carbon into compounds of smaller molecular weight to incorporate into the cell (20, 22) and CAZymes and peptidases are key enzymes for carbohydrate and protein degradation, respectively (23, 24). To account for differences in biological processes at different depths, we normalized the gene and transcript abundance of predicted secretory CAZymes and peptidases by those of all CAZyme and peptidase ORFs in metatranscriptomic and metagenomic samples (the bathypelagic zone was excluded for all transcript analysis, because no metatranscriptomes were available for Malaspina samples).The large number of CAZymes and peptidases in the database make it challenging to identify their exact binding substrates, which might negatively impact our ability to quantify the particle association of a microbial community. However, microorganisms often use biofilms to attach to the surfaces of particles (22), and thus we paired the CAZyme/peptidase analysis with a secondary analysis quantifying the abundance of biofilm-associated genes using a manually curated database of biofilm-associated ORFs (see Table S3 for the database of biofilm-associated genes).

From the *Tara* Oceans metagenomes, we identified 52,890 secretory CAZymes out of 421,080 CAZyme ORFs, and 179,932 secretory peptidases out of 1,294,210 peptidase ORFs; from the Malaspina metagenomes, we identified 7,002 secretory CAZymes in 40,415 CAZyme ORFs, and 18,014 secretory peptidases out of 67,617 peptidases (see Methods). We observed an increasing percentage of secretory CAZymes and peptidases with depth in both metagenomic and metatranscriptomic samples (**Fig. S2A-D**), which was consistent with previous findings (20). The biofilm-associated ORFs showed a similar increase in gene and transcript abundance with depth (**Fig. S2E, F**). The increase in the percentage of secretory CAZymes and peptidases with depth indicates increased extracellular enzyme activity, suggesting a shift toward a particle-associated lifestyle toward the deep ocean.

Marine particles are thought to be hotspots for horizontal gene transfer and transposase propagation (24), and the increased importance of a particle-associated lifestyle with depth in the ocean offers a promising explanation for the elevated transposase abundance in the deep ocean. In both metagenomic and metatranscriptomic samples, the abundance of transposases and the percentage of secretory CAZymes and peptidases were strongly correlated (**Fig. 2**). Given a model with only depth as predictor of the transposase abundance in a sample, the addition of secretory CAZymes and peptidases each significantly improved the accuracy of prediction (both ANOVA *F*-test *p* = 3x10^-7^, Table S3). In addition, the percentage of secretory CAZymes and peptidases each had consistent positive correlations with transposase gene and transcript abundance in each depth (**Fig. S3**). The depth independent association between particle-associated lifestyle and transposase abundance was also supported by persistent correlations between the abundance of transposase genes/transcripts with those of biofilm-associated ORFs in each depth (**Fig. S4**). Correlations in metagenomes and metatranscriptomes suggested that transposases are more abundant and more frequently transcribed in microbial communities with a greater reliance on a particle-associated lifestyle. After observing such trends at the community level, we sought to determine if these trends hold true at a genome-resolved level to support our hypothesis that the particle-associated lifestyle is a main driver for the high transposase abundance in the deep ocean.

**Fig. 2:**
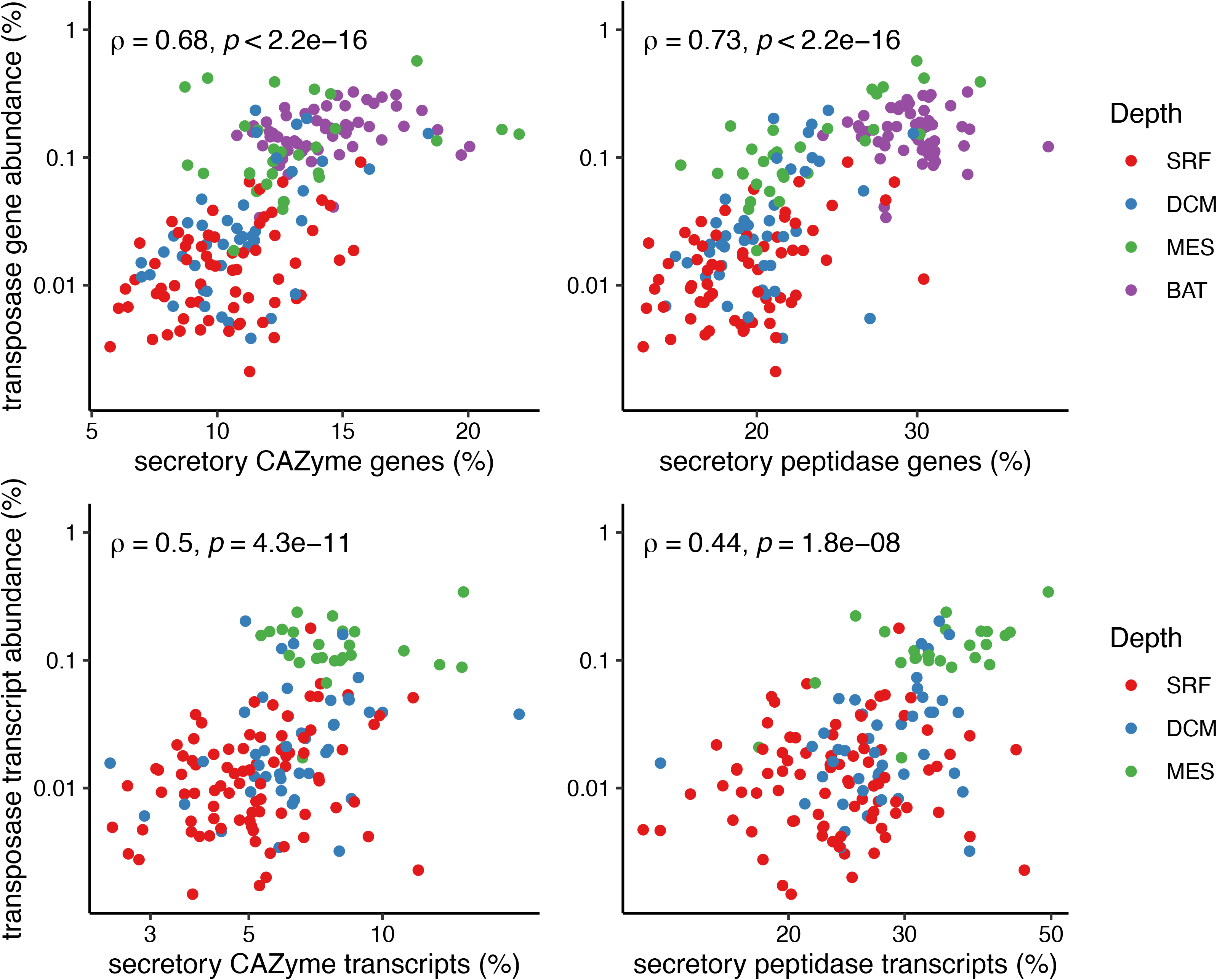
The gene potential (DNA) and transcript abundance (RNA) of transposases correlates with those of secretory CAZymes and peptidases. **A** The correlation between the abundance of transposases and secretory CAZyme and peptidases in metagenomes. **B** The same correlation in metatranscriptomes. Colors represent different ocean depths. Note that the percent transcript abundances were log-transformed.

### Genome size and taxonomy are key factors to transposase abundance in populations

To identify factors affecting transposase abundance at a genome-resolved level, we analyzed 1147 MAGs recovered from the *Tara* Oceans metagenomes (19) and 255 MAGs from the Malaspina metagenomes (18). All MAGs in our analysis had percent completeness >70%, redundancy <10%, and contained <5% of eukaryotic sequences (see Methods). The transposase abundance in MAGs was quantified by the percentage of transposase ORFs among all ORFs in the MAG (abbreviated as *%-transposase*). Similar to metagenomes, MAGs from deep waters had higher transposase abundance than MAGs from shallow waters. Specifically, MAGs from the mesopelagic and bathypelagic zones had a mean %-transposase 6 times (95% CI: 5.38 to 6.57) that of MAGs from the surface and DCM.

Given the enrichment of transposase ORFs in deep water MAGs, we ask 3 questions: 1) whether the correlation between transposase abundance and the particle-associated lifestyle persists in MAGs; 2) since larger genomes have an elevated rate of HGT, whether genome size correlates to transposase abundance in MAGs; and 3) whether specific taxa encode more transposases than others, and thus whether community composition plays a role in determining transposase abundance.

As with the metagenomes, we quantified the degree to which individual genomes were particle-associated by calculating the percentage of secretory CAZymes and peptidases out of all CAZyme and peptidase ORFs (a ratio of counts) in a MAG. This method has been verified in metagenomes separated through serial filtering techniques – MAGs assembled from samples with larger filter sizes contain a higher percentage of secretory CAZymes and peptidases in the genome compared to those filtered with smaller size fractions (25). We also independently validated this method by comparing the percentage of secretory CAZyme and peptidase ORFs in taxa generally known to be particle-associated or planktonic (**Fig. S5**). Taxonomy was analyzed at a class level, and the “complete genome size” was estimated using the genome length of a MAG divided by its estimated percent completeness. Although MAG assembly might bias against MGEs (26) and the completeness of MAGs might be underestimated for rare taxonomy groups (27, 28), we can still gain valuable insights by analyzing the distribution of transposase ORFs in MAGs of various genome sizes. To address the first question, we observed a positive correlation between the percentage of transposase ORFs and the percentage of secretory CAZymes and peptidases in MAGs (Spearman ρ = 0.208 and ρ = 0.210, respectively; both *p* < 10^-14^), thus confirming that the correlation between transposases and particle association persists on a genome-by-genome basis.

To examine the relationship between genome size and transposase abundance, we compared the estimated genome size of each MAG with the percentage of ORFs characterized as transposases within each MAG. In every depth, we observed a positive correlation between the estimated genome size and the percentage of transposase ORFs within a MAG (Spearman ρ was 0.56, 0.50, 0.46, 0.60 for surface, DCM, mesopelagic, and bathypelagic samples, respectively. All *p* < 10^-20^), confirming previous speculations that larger genomes were linked to high transposase abundance (10, 13). While deep water genomes were bigger (**Fig. 3A**), the particle-associated lifestyle correlated with greater complete genome size in every depth (**Fig. 3B**). Previous work has shown that large genomes are associated with a high fraction of transposase ORFs in isolated NCBI microbial genomes (11, 29, 30), and in a study of metagenomes from the Baltic Sea (10). The correlation between transposase abundance and genome size was previously attributed to a higher frequency of HGT in larger genomes (23, 29). The correlation between the particle-associated lifestyle and genome size has also been previously observed and may be linked to the expanded metabolic versatility of particle-associated microbial populations (25). Thus, the particle-associated lifestyle might facilitate transposase propagation by reducing the distance between microorganisms, and may lead to larger genome size through increased rates of HGT. It is worth noting that the transposase abundance in MAGs peaked when the complete genome size reached 5 Mbp, but it stabilized or declined beyond the peak (**Fig. 3C**). A previous study has also reported a decreased proportion of transposase ORFs after bacterial genomes surpass 6 Mbp in size (11). However, most (88.9%) MAGs recovered from the *Tara* Oceans and Malaspina metagenomes were under 5 Mbp, so the association between large genomes and high transposase abundance generally held true.

**Fig. 3:**
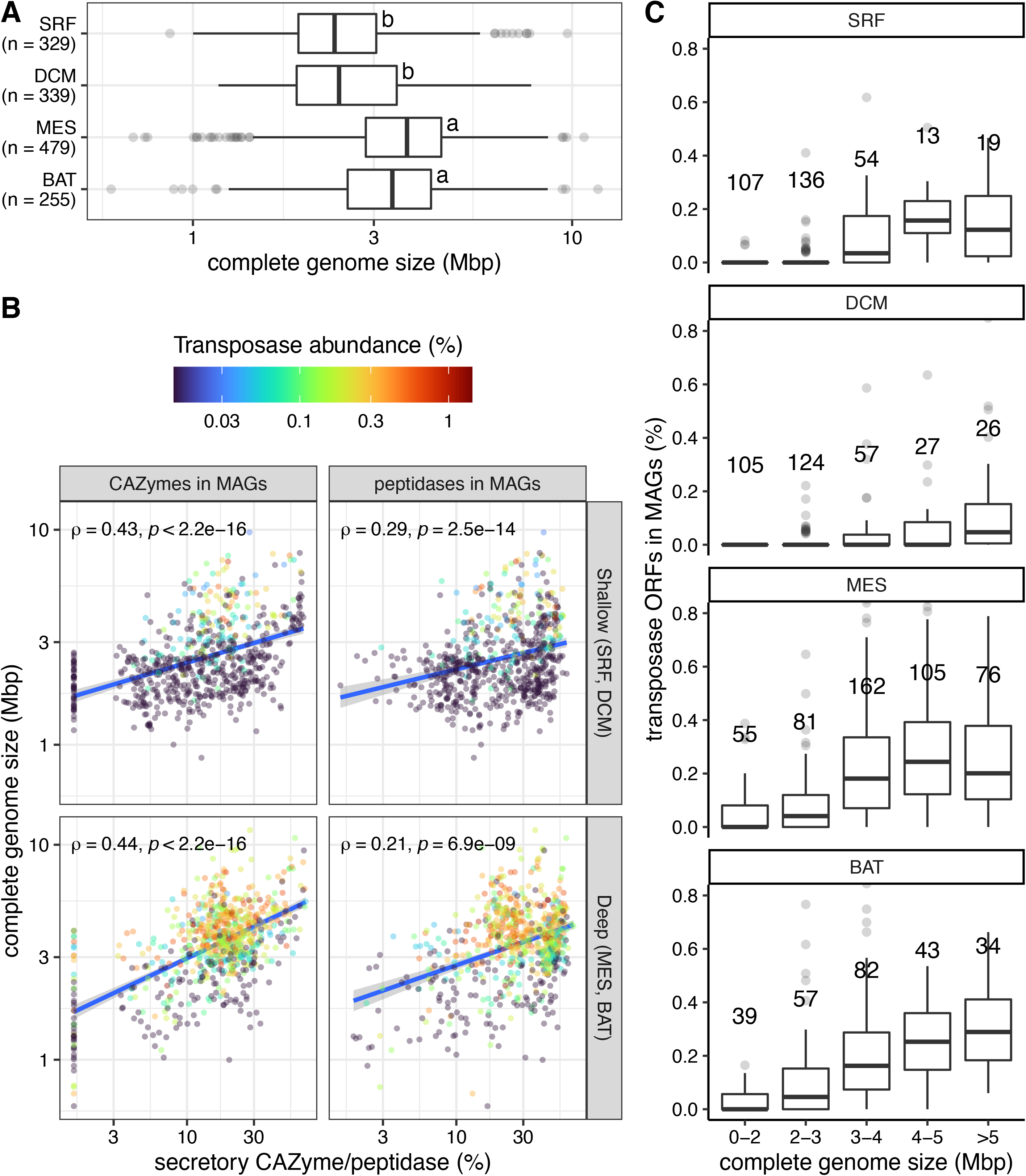
Deep-sea and particle-associated microbial populations tend to have larger genomes, which correlate with high transposase abundance in those populations. **A** The estimated complete genome size of MAGs (complete genome size = number of base pairs in a MAG / % completeness), grouped by depth. Letters on boxplots were generated from the Tukey HSD test. **B** Scatterplots showing the correlation between secretory CAZymes/peptidases and the estimated complete genome size of MAGs. **C** The transposase abundance in MAGs, grouped by depth and complete genome size. See Methods for the determination of depth of a MAG.

To address the third question, we found that taxonomic class was also a key predictor of transposase abundance in MAGs; information on the taxonomy and depth of MAGs together explained 53% of the variance in transposase abundance (**Table 1**). MAGs of the taxonomic classes *Alpha-*, *Beta-*, *Gammaproteobacteria*, and *Actinobacteria* were enriched in transposases compared to other MAGs (for each class with ≥10 MAGs, The %-transposase of MAGs in that class were compared against those of all other MAGs; Wilcox-test *p* cutoff: 0.05). MAGs of *Flavobacteria*, *Acidimicrobidae*, *novelClass_E*, and *SAR202-2* had low transposase abundance. Taxa that were enriched/low in transposases mostly matched with previous studies (10, 11), except that *Actinobacteria* had previously been reported to be low in transposase abundance.

**Table 1.** Multiple Stepwise Regression for log-transformed transposase abundance (%) of 1402 MAGs. An ANOVA test was performed on each newly added covariate.

| Explanatory Variable(s) | $p$ for $F$-test | Cumulative $R^2$ |
| --- | --- | --- |
| depth | $< 10^{-10}$ | 0.34 |
| depth + secretory CAZyme (%) | $< 10^{-10}$ | 0.37 |
| depth + secretory peptidase (%) | $< 10^{-10}$ | 0.37 |
| depth + genome size (Mbp) | $< 10^{-10}$ | 0.52 |
| depth + taxon (class) | $< 10^{-10}$ | 0.53 |
| depth + taxon + genome size + secretory CAZyme + secretory peptidase | NA | 0.62 |

The link between taxonomy and transposase abundance could partially be attributed to depth. MAGs of transposase-enriched classes were more abundant in deep waters, and MAGs from low-transposase classes were more abundant in shallow waters (2 *χ^2^*-tests, both *p* < 1x10^-6^; **Fig. S6**). Thus, although community composition is a key covariate to transposase abundance, the distribution of taxa is nonetheless linked to the depth of a microbial community.

### Relaxed selection does not explain transposase abundance in genomes

It is possible that transposases accumulate in deep ocean microbial populations as a result of genetic drift, because those populations tend to be smaller in size (31) and experience slow growth rates (32). To examine this possibility, we tested whether the strength of selection experienced by populations correlated with their transposase abundance. For each ORF in each MAG, we calculated its *pN/pS* ratio in each sample from its designated depth (*pN/pS* was only calculated for ORFs with ≥20x coverage). *pN*/pS represents the proportion of nonsynonymous mutations to the proportion of synonymous mutations, and characterizes selection at the level of the population, in contrast to *dN/dS*, which characterizes selection between individual species (33–35).

We observed that MAGs from the mesopelagic zone had a higher median *pN/pS* than MAGs from the surface and DCM, and MAGs from the bathypelagic zone had a higher median *pN/pS* than MAGs from the mesopelagic zone (both *p* < 5 x 10^-6^, **Fig. 4A**). The high median *pN/pS* of populations from the deeper layers suggested relaxed selection pressure relative to populations from shallower layers, which was in agreement with previous results (13). The general explanation for such a trend is a slower replication rate, which leads to smaller effective population sizes and a reduced selection effect in the deep ocean (12, 13). However, further work is needed to substantiate this hypothesis.

**Fig. 4:**
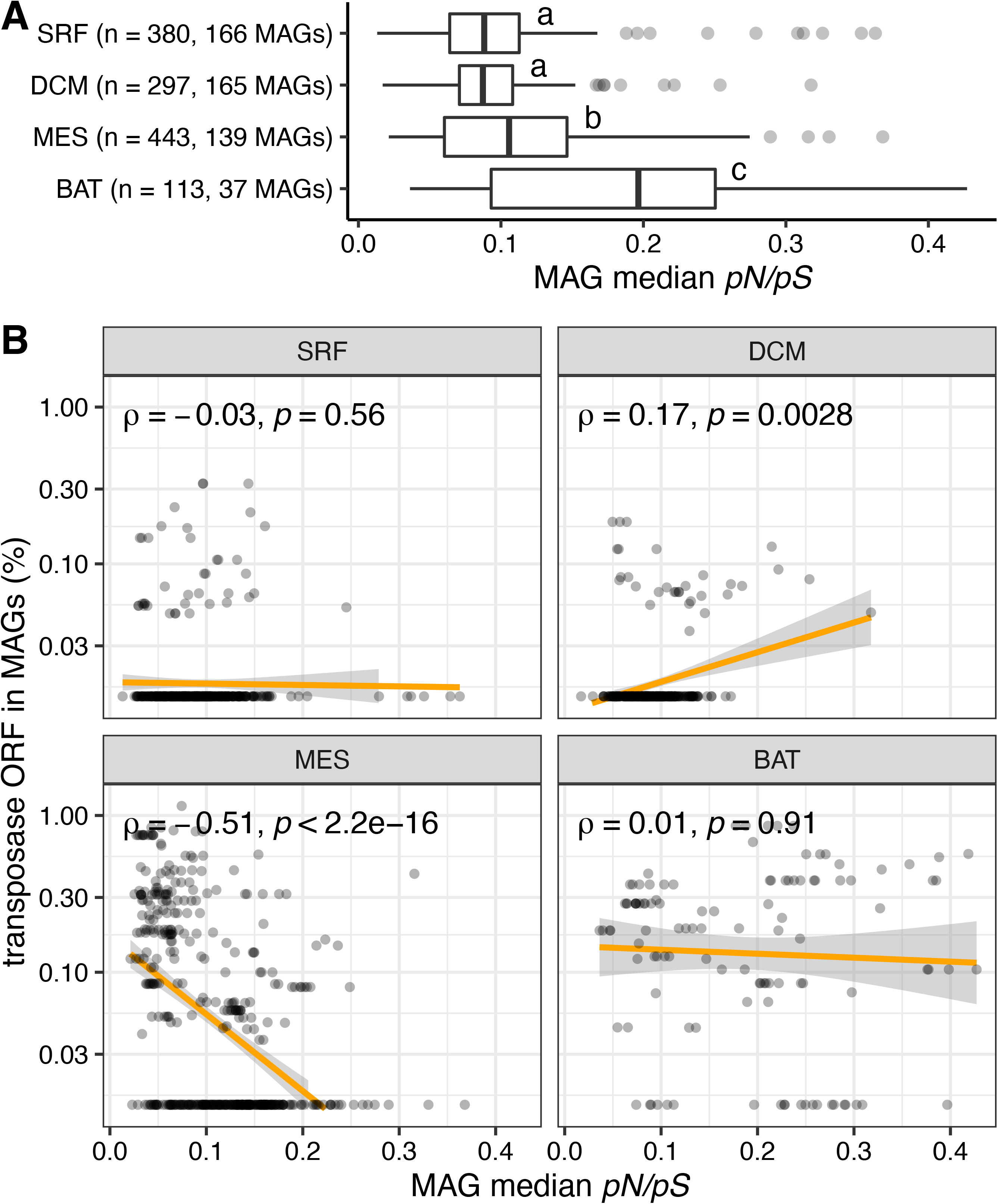
Selection pressure and transposase abundance are not correlated on a genome-resolved scale. **A** The median *pN/pS* ratio of MAGs, grouped by depth. Since *pN/pS* were calculated on a per-sample basis, a MAG might have several median *pN/pS* ratio from multiple samples. **B** The relationship between the selection pressure (median *pN/pS*) and the transposase abundance of a MAG, separated by depth. Only MAGs with ≥ 100 *pN/pS* were computed for median *pN/pS*.

Since transposases and deep-sea ORFs were both under relaxed selection, we tested whether high transposase abundance correlated with relaxed selection in a population. However, we did not observe a correlation between the median *pN/pS* of MAGs and their transposase abundance (Spearman ρ = -0.075, *p* = 0.08). Furthermore, the relationship between the selective pressure experienced by a population and its transposase abundance was inconsistent across different depths (**Fig. 4B**). It has been previously suggested that small populations would experience relaxed selection against mobile genetic elements (36, 37). However, our analyses show that although relaxed selection and transposase enrichment co-occur in deep oceans, we did not observe this trend on a genome-by-genome basis, suggesting that there is not a direct connection between transposase abundance and relaxed selection at a genome-resolved scale, and thus within individual microbial populations.

### Transposons/integrons carry a high proportion of ORFs related to defense mechanisms

One important question regarding high transposase abundance in the deep sea is whether these transposases are neutral or deleterious, or whether they perform important functions in the genomes encoding them. In some cases, transposases are deleterious and proliferate as a result of genetic drift (38), but transposases can also benefit the host by introducing functional and regulatory genes (7, 24, 39, 40).

Transposases mediate the migration and integration of cassette sequences into host genomes; a transposon or integron is the combination of a transposase and its cassettes. Thus, cassette sequences are functionally similar to auxiliary metabolic genes (AMGs) (41), in that they are carried by a selfish MGE (or viruses in the case of AMGs) to increase the fitness of the host and therefore increase the fitness of the MGE.

To determine whether transposons in marine habitats carry advantageous genes, we began by querying the functional categories of cassette sequences. The software package Integron Finder (11) was used to locate cassettes on contigs; the program searches for the two palindromic flanking sites necessary for transposase-mediated recombination as well as a nearby transposase/integrase. We treated transposons and integrons as equivalent, due to the sequence and function similarity between integrases and transposases (11). We found 8,519 cassette ORFs from the co-assembled Malaspina (18) and *Tara* Oceans (17, 19) metagenomes (see Methods). Only 27% of the cassette sequences were assigned with clusters of orthologous genes (COG) annotations, which was a lower annotation rate compared to other ORFs in the *Tara* Oceans (55-60%) and Malaspina (68%) metagenomes.

Compared to the rest of the metagenome, cassettes were enriched in ORFs of the COG categories “replication, recombination, and repair”, “defense mechanisms”, and “mobilome: prophages, transposons” (**Fig. 5A**). Transposons/integrons are expected to be enriched in ORFs related to the mobilome and replication/recombination. The prevalence of defense mechanism genes in cassettes was noteworthy: defense mechanisms accounted for 14.3% of known function calls in cassettes, but only accounted for 2.47% of known function calls in non-cassette ORFs (*known* was defined as all functional groups except “function unknown” and “general function prediction”). Out of the 276 defense mechanism cassettes, 123 of them encoded toxin/antitoxin genes. Since transposases are more pervasive on particles, toxin and defense genes they carry would be useful in competition and protection on crowded particles (42–45). Moreover, the percentage of secretory CAZymes and peptidases in metagenomes both correlated with the abundance of defense mechanism ORFs (**Fig. 5B**). Similarly, we also found a strong correlation between the gene abundance of defense mechanism ORFs and that of biofilm-associated ORFs (**Fig. S7**). Thus, if defense mechanism genes in the deep ocean were more highly expressed, this would substantiate the hypothesis that deep ocean microorganisms benefit from novel genes introduced by integrons/transposons.

**Fig. 5:**
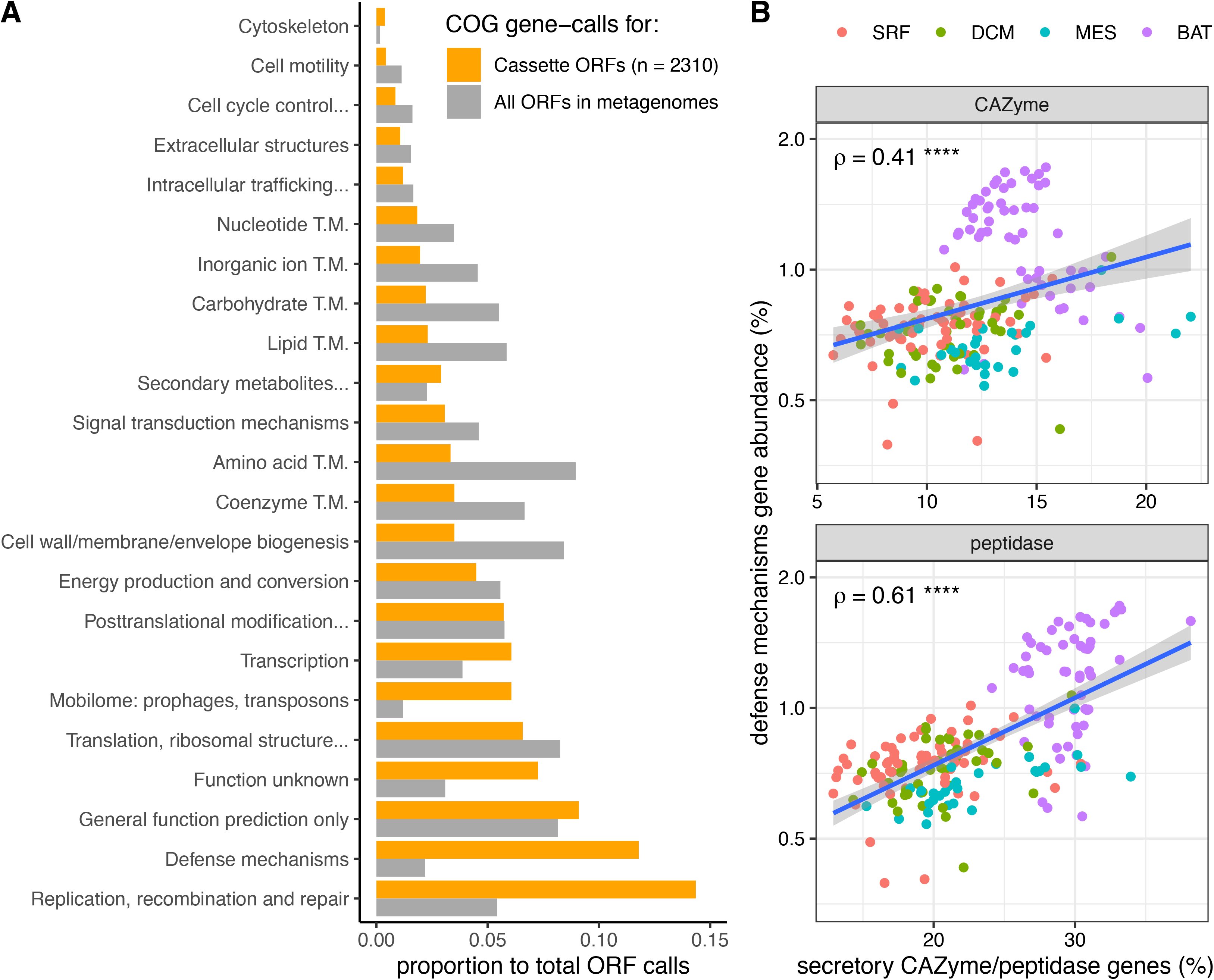
Cassette sequences include high proportions of defense mechanism genes, which are more abundant in microbial communities that rely heavily on a particle-associated lifestyle. **A** The distribution of COG functional categories in cassette and non-cassette metagenome ORFs. *TM: transport and metabolism. 1951 cassette ORFs with COG function calls were identified from the 10 co-assembled *Tara* metagenomes, and 359 from the co-assembled Malaspina metagenome. 5 million ORFs with COG function calls were then sampled from the *Tara* Oceans and Malaspina metagenomes according to the ratio of identified cassettes described above. **B** The correlation between the abundance of secretory CAZyme/peptidase ORFs and defense mechanism ORFs in metagenomes. Samples from different depths are distinguished by different colors. The Spearman’s correlation coefficients, ρ, are shown on top left. **** indicates that *p* < 0.0001.

### ORFs related to defense mechanisms, secretory CAZymes, and biofilm-associated genes have higher expression in the deep sea

To determine whether defense mechanism genes are more highly expressed and how this correlates to expression of particle-associated genes, we calculated their RNA/DNA ratios (defined as the transcript abundance of a target gene divided by its gene abundance) of secretory CAZyme, secretory peptidase, defense mechanism, and transposase ORFs in each *Tara* Oceans sample with a paired metagenome and metatranscriptome (n = 94). Malaspina samples were excluded from this analysis, because no metatranscriptomes were sequenced.

The ORFs related to secretory CAZymes and defense mechanisms had greater RNA/DNA ratios in the mesopelagic zone than in the surface and DCM (**Fig. 6**), demonstrating a greater need for these genes as deep ocean microbial communities switch toward a particle-associated lifestyle. The RNA/DNA ratio of biofilm-associated ORFs also supported this switch to particle-associated lifestyle (**Fig. S8**). In contrast, although transposases were more abundant in the mesopelagic zone (**Fig. S9**), their RNA/DNA ratios were similar across all depths (**Fig. 6**). A possible explanation is that although transposases are not upregulated, they sometimes carry beneficial cassette genes, which are increasingly expressed in deep-sea microbial genomes.

**Fig. 6:**
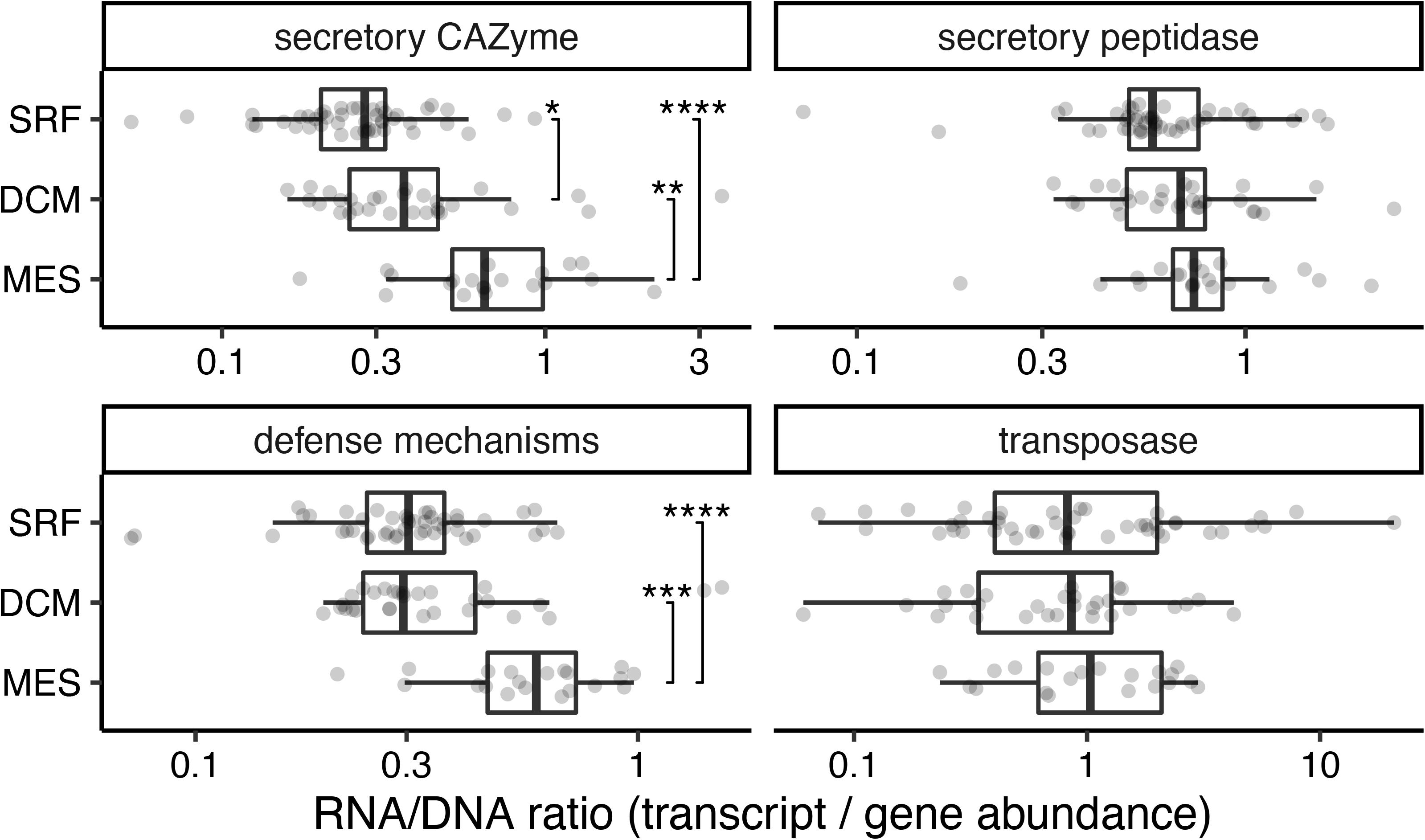
Particle-associated and defense mechanism ORFs are more highly expressed in deeper waters, but transposases are not. Log-transformed RNA/DNA ratios of target genes in each sample, separated by depth. * *p* < 0.05, ** *p* < 0.01, *** *p* < 0.001.

## Conclusion

Our analysis provides insights into the factors driving the increasing abundance of transposases with ocean depth, highlighting the interplay between the particle-associated lifestyle, expanded genome size, and selection for defense mechanism genes (**Fig. 7**). We hypothesize that as microbial communities shift from a predominantly planktonic to particle-associated lifestyle from the surface to the bathypelagic zone, microbial communities become more densely packed on particles, leading to rampant transposase spread and more intense resource competition. Additionally, particle-associated microorganisms tend to have larger genomes, which are associated with high transposase abundance. These high abundances of mobile genetic elements actively shape the ecology of particle-associated microbial communities by introducing novel genes, with cassette sequences being particularly enriched in defense mechanism genes. These genes are highly expressed in the deep ocean, offering competitive advantages in the competitive particle-associated environment (42, 44).

**Fig. 7:**
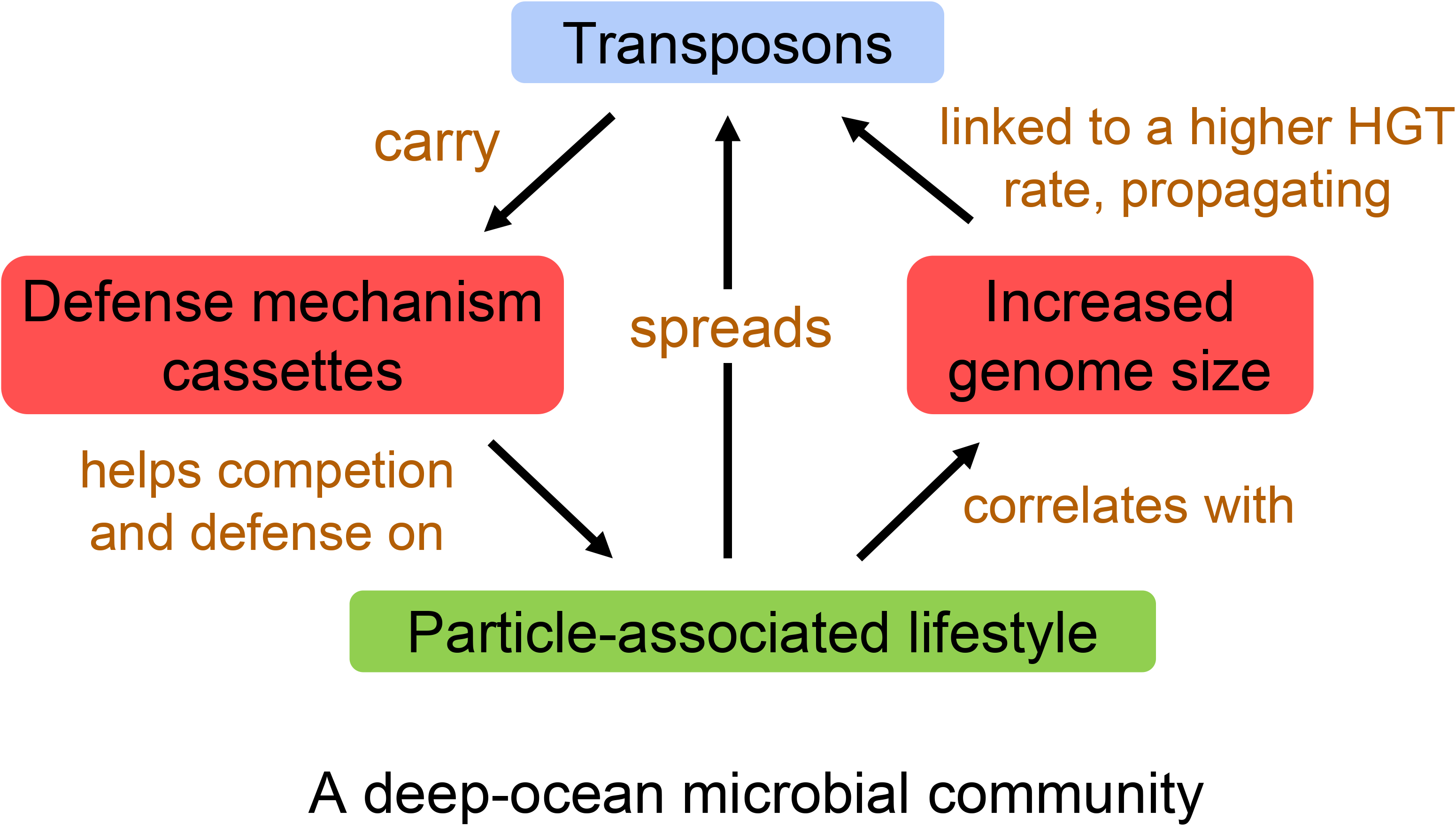
Linking a particle-associated lifestyle to the high abundance of transposons in the deep ocean.

When interpreting these results, it is important to consider certain caveats. Our analysis focused on pelagic ocean water samples to minimize potential confounding variables, and thus these observed trends may not apply to regions with unique characteristics, such as Arctic sea ice (47) and deep-sea hydrothermal vents (15). Moreover, as our results are based on correlative analyses, further experimental evidence is needed to establish causation and to identify mechanistic relationships between these variables.

Understanding these trends in mobile genetic element abundance is important for understanding gene flow, competition, and other ecological characteristics of the dark ocean, one of the largest habitats on Earth. Our results show a strong association between transposase abundance and a particle-associated lifestyle, suggesting that transposons may enable microbial lineages to compete in crowded biofilm-associated habitats. We did not find any correlation between the strength of selection and transposase abundance at the genome level, indicating that relaxed selection pressure is not the primary driver for the high transposase abundance in the deep ocean. Overall, our results suggest an emerging understanding of the ocean as a stratified system in which the deep ocean acts as a gene sharing highway, fostering networks of gene exchange, particularly on particles. This results in large genomes with many defense-oriented genes in the deep-sea microbial communities, contrasting with the streamlined and specialized genomes that dominate the surface oceans. Future experimental studies should establish mechanistic connections between transposase gene cassette contents and microbial activity, particularly in planktonic and particle-associated communities.

## Materials and Methods

### Analysis of the Ocean Microbial Reference Catalog v2 (OM-RGC.v2) from the *Tara* Oceans Project

OM-RGC.v2 contained 47 million non-redundant open reading frames (ORFs) (16, 17), and the coverage of each ORF in every metagenomic (n = 138) and metatranscriptomic (n = 152) sample (<u>website</u>) (16). Samples from the Arctic Ocean were excluded. We identified transposases and biofilm-associated ORFs in OM-RGC.v2 through TBLASTN (e-value < 10^-5^). The transposase database contained “transposase” and “integrase” genes from the Pfam database (48) (Table S1). We identified ORFs of the “defense mechanisms” category through COG annotations in OM-RGC.v2. Finally, the coverage data in OM-RGC.v2 was used to determine the relative abundance of a target gene in a sample.

### Acquisition of global ocean metagenomes

Metagenomic reads of the *Tara* Oceans Project and Malaspina Expedition were downloaded from the European Nucleotide Archive (ENA) under study accessions PRJEB402 and PRJEB44456, respectively. Methods for sample collection and Illumina sequencing were described by Sunagawa *et al.* (49) for the *Tara* Oceans Project and Acinas *et al.* (18) for the Malaspina Expedition. For the assembled contigs used as references for mapping of the *Tara* Oceans metagenomes, we used 10 existing co-assembled metagenomes from the 10 oceanic provinces (19). For the reference contigs of the Malaspina metagenomes, we downloaded the co-assembled metagenome of all 58 samples (18). Samples from the *Tara* Oceans were filtered through 1.6- or 3-μm filters and collected on 0.22-μm filters (49), whereas samples from the Malaspina expedition were collected from the 0.2 – 0.8 μm and 0.8 – 5 μm size fractions (18). We used EukDetect (50) to confirm that samples collected with a larger size fraction did not have a higher contribution from eukaryotic reads (see Supplementary Text).

### Mapping and calculating *pN/pS* ratios

To assess the strength of selection of target ORFs, we calculated the *pN/pS* ratio (the proportion of nonsynonymous mutations over the proportion of synonymous mutations) for all ORFs. To do this, we mapped the metagenomic reads against the assembled contigs for each metagenome. We mapped the raw reads of each of the *Tara* Oceans metagenomes to the co-assembly from their corresponding provinces, and all raw reads of each of the Malaspina metagenomes to one co-assembled metagenome using Bowtie2 (51) (v2.2.9; paired-end alignment with default parameters). We calculated *pN/pS* using anvi’o (52) with the script “anvi-script-calculate-pn-ps-ratio” with the “--min-coverage” flag set to 20. The *pN/pS* ratios were calculated on a sample-per-sample basis, so an ORF from a co-assembled metagenome might have multiple *pN/pS* ratios from different samples.

### Prediction of secretory CAZyme and peptidase ORFs

We adapted the methods employed in Zhao et al (20) to assess whether microbial communities were largely planktonic or particle-associated by quantifying the relative abundance of secretory CAZymes and peptidases. Predicted CAZymes and peptidases were annotated using DIAMOND (2.0.15) (53) BLASTp (e-value <10^−10^) to search against the dbCAN (54) and MEROPS (55) databases, respectively. SignalP (5.0) (56) was used to identify signal peptides. We used the Gram-positive mode for ORFs affiliated with *Actinobacteria* and *Firmicutes*, Gram-negative mode for other bacterial phylum, and Archaea mode for ORFs affiliated with the *Archaea* domain. Only ORFs in the *Bacteria* or *Archaea* domains were included in the analysis. For MAGs, counts of secretory CAZymes and peptidases were used instead of abundance; validation based on predicted lifestyles of known taxa confirmed that this gave an accurate prediction of lifestyle (see Supplementary Text).

### MAG Selection

To conduct genome-resolved analyses, we selected 1147 out of 2631 MAGs that had previously been recovered from the Tara Ocean metagenomes (19) and 255 out of 317 MAGs recovered from the Malaspina metagenomes (18). All selected MAGs had <5% of eukaryotic sequences (in base pairs, Tiara (*57*) was used to identify putative eukaryotic contigs), completeness > 70%, and redundancy < 10% (both criteria were calculated with CheckM (58)).

### Determination of MAG’s depth and lifestyles

MAGs that originated from co-assemblies in a specific province in the *Tara* Oceans dataset (*e.g.*, Red Sea), as in Tully *et al.* (19), were used to recruit reads from all *Tara* Oceans samples (n = 180) using Bowtie2 (default parameters). Each province was performed separately. SAM files were converted to BAM files using samtools (59) and used to determine RPKM (reads per kilobase pair MAG per Million pair metagenome) as in Graham et al. (60).

To assign the depth origin of a MAG from the *Tara* Oceans metagenomes, we summed up its RPKM in surface, DCM, and mesopelagic samples (3 groups, separately). The layer with the highest sum RPKM was the depth of that MAG. All Malaspina MAGs belonged to the bathypelagic layer.

### Integron and cassette sequences detection

We used Integron Finder (v2) (11) to identify cassette sequences from the *Tara* Ocean and Malaspina metagenomes. Integron Finder uses HMMER to locate the integron-integrase *intI*, which is conserved for most integrons. Then, cassette sequences are identified with the near-palindromic flaking regions. anvi’o (52) was used to identify and annotate ORFs on metagenomic contigs using the script ‘anvi-run-ncbi-cogs.’ If the anvi’o ORF calls from Prodigal (61) were within 100bps (start and stop) of the Integron Finder’s cassette calls, the anvi’o COG annotations were used for those cassette ORFs.

### Statistical analysis

We used the Wilcoxon signed-rank test to compare the differences between means of two numeric variables, and the Spearman’s rank correlation was used to determine the association between two numeric variables. Multiple hypothesis *p-*values were adjusted using the Benjamini–Hochberg procedure. All linear regressions had a normal distribution of residuals (Shapiro–Wilk test).

Statistical significance was assumed if *p* < 0.05. All statistical analyses were performed using base R packages (v4.1.3).

## Code and data availability

All Python, R scripts, code explanation, and raw data for analysis are publicly accessible on GitHub at https://github.com/carleton-spacehogs/transposase-deep-ocean.

## Acknowledgments

We would like to thank Dr. Shinichi Sunagawa for kindly providing information about the *Tara* Oceans datasets and the Ocean Microbial Reference Catalog, and Dr. Silvia G. Acinas for providing information about the Malaspina Deep Ocean dataset. Dr. Murat Eren provided help with anvi’o, and Mike Tie provided assistance with server administration. Funding for JZ and OK was provided by grants from the Towsley Endowment at Carleton College, and funding for TdRH was provided by the Summer Science Fellows program at Carleton College.

## References

1. Frost LS, Leplae R, Summers AO, Toussaint A. 2005. Mobile genetic elements: the agents of open source evolution. Nat Rev Microbiol 3:722–732.

2. Transposons. http://www.nature.com/scitable/topicpage/transposons-the-jumping-genes-518. Retrieved 30 November 2021.

3. Muñoz-López M, García-Pérez JL. 2010. DNA transposons: nature and applications in genomics. Curr Genomics 11:115–128.

4. Collis C M, Hall R M. 1995. Expression of antibiotic resistance genes in the integrated cassettes of integrons. Antimicrob Agents Chemother 39:155–162.

5. Rankin DJ, Rocha EPC, Brown SP. 2011. What traits are carried on mobile genetic elements, and why? Heredity 106:1–10.

6. Hacker J, Carniel E. 2001. Ecological fitness, genomic islands and bacterial pathogenicity. A Darwinian view of the evolution of microbes. EMBO Rep 2:376–381.

7. Jones JM, Grinberg I, Eldar A, Grossman AD. 2021. A mobile genetic element increases bacterial host fitness by manipulating development. Elife 10:e65924.

8. Casacuberta E, González J. 2013. The impact of transposable elements in environmental adaptation. Mol Ecol 22:1503–1517.

9. Aziz RK, Breitbart M, Edwards RA. 2010. Transposases are the most abundant, most ubiquitous genes in nature. Nucleic Acids Res 38:4207–4217.

10. Vigil-Stenman T, Ininbergs K, Bergman B, Ekman M. 2017. High abundance and expression of transposases in bacteria from the Baltic Sea. ISME J 11:2611–2623.

11. Cury J, Jové T, Touchon M, Néron B, Rocha EP. 2016. Identification and analysis of integrons and cassette arrays in bacterial genomes. Nucleic Acids Res 44:4539–4550.

12. DeLong EF, Preston CM, Mincer T, Rich V, Hallam SJ, Frigaard N-U, Martinez A, Sullivan MB, Edwards R, Brito BR, Chisholm SW, Karl DM. 2006. Community Genomics Among Stratified Microbial Assemblages in the Ocean’s Interior. Science 311:496–503.

13. Konstantinidis KT, Braff J, Karl DM, DeLong EF. 2009. Comparative metagenomic analysis of a microbial community residing at a depth of 4,000 meters at station ALOHA in the North Pacific subtropical gyre. Appl Environ Microbiol 75:5345–5355.

14. Worden AZ, Cuvelier ML, Bartlett DH. 2006. In-depth analyses of marine microbial community genomics. Trends Microbiol 14:331–336.

15. Brazelton WJ, Baross JA. 2009. Abundant transposases encoded by the metagenome of a hydrothermal chimney biofilm. ISME J 3:1420–1424.

16. Salazar G, Paoli L, Alberti A, Huerta-Cepas J, Ruscheweyh H-J, Cuenca M, Field CM, Coelho LP, Cruaud C, Engelen S, Gregory AC, Labadie K, Marec C, Pelletier E, Royo-Llonch M, Roux S, Sánchez P, Uehara H, Zayed AA, Zeller G, Carmichael M, Dimier C, Ferland J, Kandels S, Picheral M, Pisarev S, Poulain J, Tara Oceans Coordinators, Acinas SG, Babin M, Bork P, Bowler C, de Vargas C, Guidi L, Hingamp P, Iudicone D, Karp-Boss L, Karsenti E, Ogata H, Pesant S, Speich S, Sullivan MB, Wincker P, Sunagawa S. 2019. Gene Expression Changes and Community Turnover Differentially Shape the Global Ocean Metatranscriptome. Cell 179:1068–1083.e21.

17. Sunagawa S, Coelho LP, Chaffron S, Kultima JR, Labadie K, Salazar G, Djahanschiri B, Zeller G, Mende DR, Alberti A, Cornejo-Castillo FM, Costea PI, Cruaud C, d’Ovidio F, Engelen S, Ferrera I, Gasol JM, Guidi L, Hildebrand F, Kokoszka F, Lepoivre C, Lima-Mendez G, Poulain J, Poulos BT, Royo-Llonch M, Sarmento H, Vieira-Silva S, Dimier C, Picheral M, Searson S, Kandels-Lewis S, Null N, Bowler C, de Vargas C, Gorsky G, Grimsley N, Hingamp P, Iudicone D, Jaillon O, Not F, Ogata H, Pesant S, Speich S, Stemmann L, Sullivan MB, Weissenbach J, Wincker P, Karsenti E, Raes J, Acinas SG, Bork P, Boss E, Bowler C, Follows M, Karp-Boss L, Krzic U, Reynaud EG, Sardet C, Sieracki M, Velayoudon D. 2015. Structure and function of the global ocean microbiome. Science 348:1261359.

18. Acinas SG, Sánchez P, Salazar G, Cornejo-Castillo FM, Sebastián M, Logares R, Royo-Llonch M, Paoli L, Sunagawa S, Hingamp P, Ogata H, Lima-Mendez G, Roux S, González JM, Arrieta JM, Alam IS, Kamau A, Bowler C, Raes J, Pesant S, Bork P, Agustí S, Gojobori T, Vaqué D, Sullivan MB, Pedrós-Alió C, Massana R, Duarte CM, Gasol JM. 2021. Deep ocean metagenomes provide insight into the metabolic architecture of bathypelagic microbial communities. Communications Biology 4:1–15.

19. Tully BJ, Graham ED, Heidelberg JF. 2018. The reconstruction of 2,631 draft metagenome-assembled genomes from the global oceans. Scientific Data 5:1–8.

20. Zhao Z, Baltar F, Herndl GJ. 2020. Linking extracellular enzymes to phylogeny indicates a predominantly particle-associated lifestyle of deep-sea prokaryotes. Sci Adv 6:eaaz4354.

21. Ganesh S, Parris DJ, DeLong EF, Stewart FJ. 2014. Metagenomic analysis of size-fractionated picoplankton in a marine oxygen minimum zone. ISME J 8:187–211.

22. Tuson HH, Weibel DB. 2013. Bacteria-surface interactions. Soft Matter 9:4368–4380.

23. Cordero OX, Hogeweg P. 2009. The impact of long-distance horizontal gene transfer on prokaryotic genome size. Proc Natl Acad Sci U S A 106:21748–21753.

24. Stewart FJ. 2013. Where the genes flow. Nat Geosci 6:688–690.

25. Leu AO, Eppley JM, Burger A, DeLong EF. 2022. Diverse Genomic Traits Differentiate Sinking-Particle-Associated versus Free-Living Microbes throughout the Oligotrophic Open Ocean Water Column. MBio 13:e0156922.

26. Meziti A, Rodriguez-R LM, Hatt JK, Peña-Gonzalez A, Levy K, Konstantinidis KT. 2021. The Reliability of Metagenome-Assembled Genomes (MAGs) in Representing Natural Populations: Insights from Comparing MAGs against Isolate Genomes Derived from the Same Fecal Sample. Appl Environ Microbiol 87.

27. Garcia SL, Buck M, McMahon KD, Grossart H-P, Eiler A, Warnecke F. 2015. Auxotrophy and intrapopulation complementary in the “interactome” of a cultivated freshwater model community. Mol Ecol 24:4449–4459.

28. Rodríguez-Gijón A, Nuy JK, Mehrshad M, Buck M, Schulz F, Woyke T, Garcia SL. 2021. A Genomic Perspective Across Earth’s Microbiomes Reveals That Genome Size in Archaea and Bacteria Is Linked to Ecosystem Type and Trophic Strategy. Front Microbiol 12:761869.

29. Iranzo J, Gómez MJ, López de Saro FJ, Manrubia S. 2014. Large-scale genomic analysis suggests a neutral punctuated dynamics of transposable elements in bacterial genomes. PLoS Comput Biol 10:e1003680.

30. Touchon M, Rocha EPC. 2007. Causes of Insertion Sequences Abundance in Prokaryotic Genomes. Mol Biol Evol 24:969–981.

31. Costello MJ, Chaudhary C. 2017. Marine Biodiversity, Biogeography, Deep-Sea Gradients, and Conservation. Curr Biol 27:R511–R527.

32. Sogin ML, Morrison HG, Huber JA, Welch DM, Huse SM, Neal PR, Arrieta JM, Herndl GJ. 2006. Microbial diversity in the deep sea and the underexplored “rare biosphere.” Proc Natl Acad Sci U S A 103:12115–12120.

33. Simmons SL, Dibartolo G, Denef VJ, Goltsman DSA, Thelen MP, Banfield JF. 2008. Population genomic analysis of strain variation in Leptospirillum group II bacteria involved in acid mine drainage formation. PLoS Biol 6:e177.

34. McDonald JH, Kreitman M. 1991. Adaptive protein evolution at the Adh locus in Drosophila. Nature 351:652–654.

35. Schloissnig S, Arumugam M, Sunagawa S, Mitreva M, Tap J, Zhu A, Waller A, Mende DR, Kultima JR, Martin J, Kota K, Sunyaev SR, Weinstock GM, Bork P. 2013. Genomic variation landscape of the human gut microbiome. Nature 493:45–50.

36. Parkhill J, Sebaihia M, Preston A, Murphy LD, Thomson N, Harris DE, Holden MTG, Churcher CM, Bentley SD, Mungall KL, Cerdeño-Tárraga AM, Temple L, James K, Harris B, Quail MA, Achtman M, Atkin R, Baker S, Basham D, Bason N, Cherevach I, Chillingworth T, Collins M, Cronin A, Davis P, Doggett J, Feltwell T, Goble A, Hamlin N, Hauser H, Holroyd S, Jagels K, Leather S, Moule S, Norberczak H, O’Neil S, Ormond D, Price C, Rabbinowitsch E, Rutter S, Sanders M, Saunders D, Seeger K, Sharp S, Simmonds M, Skelton J, Squares R, Squares S, Stevens K, Unwin L, Whitehead S, Barrell BG, Maskell DJ. 2003. Comparative analysis of the genome sequences of Bordetella pertussis, Bordetella parapertussis and Bordetella bronchiseptica. Nat Genet 35:32–40.

37. Moran NA, Plague GR. 2004. Genomic changes following host restriction in bacteria. Curr Opin Genet Dev 14:627–633.

38. Escobar-Páramo P, Ghosh S, DiRuggiero J. 2005. Evidence for Genetic Drift in the Diversification of a Geographically Isolated Population of the Hyperthermophilic Archaeon Pyrococcus. Molecular Biology and Evolution https://doi.org/10.1093/molbev/msi227.

39. Flynn KJ, Swanson MS. 2014. Integrative conjugative element ICE-βox confers oxidative stress resistance to Legionella pneumophila in vitro and in macrophages. MBio 5:e01091–14.

40. Sullivan JT, Ronson CW. 1998. Evolution of rhizobia by acquisition of a 500-kb symbiosis island that integrates into a phe-tRNA gene. Proc Natl Acad Sci U S A 95:5145–5149.

41. Breitbart M, Bonnain C, Malki K, Sawaya NA. 2018. Phage puppet masters of the marine microbial realm. Nat Microbiol 3:754–766.

42. Nadell CD, Drescher K, Foster KR. 2016. Spatial structure, cooperation and competition in biofilms. Nat Rev Microbiol 14:589–600.

43. Basler M, Ho BT, Mekalanos JJ. 2013. Tit-for-tat: type VI secretion system counterattack during bacterial cell-cell interactions. Cell 152:884–894.

44. Hayes CS, Aoki SK, Low DA. 2010. Bacterial contact-dependent delivery systems. Annu Rev Genet 44:71–90.

45. Ho BT, Dong TG, Mekalanos JJ. 2014. A view to a kill: the bacterial type VI secretion system. Cell Host Microbe 15:9–21.

46. Mestre M, Ruiz-González C, Logares R, Duarte CM, Gasol JM, Sala MM. 2018. Sinking particles promote vertical connectivity in the ocean microbiome. Proc Natl Acad Sci U S A 115:E6799–E6807.

47. Rapp JZ, Sullivan MB, Deming JW. 2021. Divergent Genomic Adaptations in the Microbiomes of Arctic Subzero Sea-Ice and Cryopeg Brines. Front Microbiol 12:701186.

48. Mistry J, Chuguransky S, Williams L, Qureshi M, Salazar GA, Sonnhammer ELL, Tosatto SCE, Paladin L, Raj S, Richardson LJ, Finn RD, Bateman A. 2021. Pfam: The protein families database in 2021. Nucleic Acids Res 49:D412–D419.

49. Sunagawa S, Acinas SG, Bork P, Bowler C, Tara Oceans Coordinators, Eveillard D, Gorsky G, Guidi L, Iudicone D, Karsenti E, Lombard F, Ogata H, Pesant S, Sullivan MB, Wincker P, de Vargas C. 2020. Tara Oceans: towards global ocean ecosystems biology. Nat Rev Microbiol 18:428–445.

50. Lind AL, Pollard KS. 2021. Accurate and sensitive detection of microbial eukaryotes from whole metagenome shotgun sequencing. Microbiome.

51. Langmead B, Salzberg SL. 2012. Fast gapped-read alignment with Bowtie 2. Nat Methods 9:357–359.

52. Eren AM, Kiefl E, Shaiber A, Veseli I, Miller SE, Schechter MS, Fink I, Pan JN, Yousef M, Fogarty EC, Trigodet F, Watson AR, Esen ÖC, Moore RM, Clayssen Q, Lee MD, Kivenson V, Graham ED, Merrill BD, Karkman A, Blankenberg D, Eppley JM, Sjödin A, Scott JJ, Vázquez-Campos X, McKay LJ, McDaniel EA, Stevens SLR, Anderson RE, Fuessel J, Fernandez-Guerra A, Maignien L, Delmont TO, Willis AD. 2021. Community-led, integrated, reproducible multi-omics with anvi’o. Nat Microbiol 6:3–6.

53. Buchfink B, Reuter K, Drost H-G. 2021. Sensitive protein alignments at tree-of-life scale using DIAMOND. Nat Methods 18:366–368.

54. Huang L, Zhang H, Wu P, Entwistle S, Li X, Yohe T, Yi H, Yang Z, Yin Y. 2018. dbCAN-seq: a database of carbohydrate-active enzyme (CAZyme) sequence and annotation. Nucleic Acids Res 46:D516–D521.

55. Rawlings ND, Barrett AJ, Thomas PD, Huang X, Bateman A, Finn RD. 2018. The MEROPS database of proteolytic enzymes, their substrates and inhibitors in 2017 and a comparison with peptidases in the PANTHER database. Nucleic Acids Res 46:D624–D632.

56. Almagro Armenteros JJ, Tsirigos KD, Sønderby CK, Petersen TN, Winther O, Brunak S, von Heijne G, Nielsen H. 2019. SignalP 5.0 improves signal peptide predictions using deep neural networks. Nat Biotechnol 37:420–423.

57. Karlicki M, Antonowicz S, Karnkowska A. 2022. Tiara: deep learning-based classification system for eukaryotic sequences. Bioinformatics 38:344–350.

58. Parks DH, Imelfort M, Skennerton CT, Hugenholtz P, Tyson GW. 2015. CheckM: assessing the quality of microbial genomes recovered from isolates, single cells, and metagenomes. Genome Res 25:1043–1055.

59. Li H, Handsaker B, Wysoker A, Fennell T, Ruan J, Homer N, Marth G, Abecasis G, Durbin R, 1000 Genome Project Data Processing Subgroup. 2009. The Sequence Alignment/Map format and SAMtools. Bioinformatics 25:2078–2079.

60. Graham ED, Heidelberg JF, Tully BJ. 2018. Potential for primary productivity in a globally-distributed bacterial phototroph. ISME J 12:1861–1866.

61. Hyatt D, Chen G-L, Locascio PF, Land ML, Larimer FW, Hauser LJ. 2010. Prodigal: prokaryotic gene recognition and translation initiation site identification. BMC Bioinformatics 11:119.

