## Supplementary Information for "Increasing transposase abundance with ocean depth correlates with a particle-associated lifestyle"

### High transposase abundance in the deep ocean is linked to a particle-associated lifestyle

This PDF file includes:

Tables S1 to S4 (Table S1, S3 are additional CSV files)

Figures S1 to S6

Supplementary Methods

**Supplementary Tables:**

**Table S1**. CSV file containing a database of transposase genes.

**Table S2**. Comparison of transposase abundance (calculated as the percentage of metagenomic reads to total reads in a sample) in the shallow and deep water layers, separated by oceanic regions. Shallow waters: samples from the surface (SRF) and deep chlorophyll maximum (DCM). Deep waters: samples from the mesopelagic (MES) and bathypelagic (BAT) zone. Six 2-tailed *t*-tests were performed on log-transformed transposase abundance. *** $P < 0.001$, **** $P < 0.0001$.

| **Region** | **Median Transposase Abundance (%)** | | **Sample Size** | | **Adjusted *P*** |
|---|---|---|---|---|---|
| | Shallow waters | Deep waters | Shallow | Deep | |
| Indian Ocean | 0.0111 | 0.1437 | 21 | 19 | **** |
| North Atlantic | 0.0111 | 0.1552 | 16 | 20 | **** |
| North Pacific | 0.0224 | 0.1542 | 11 | 15 | *** |
| South Atlantic | 0.0329 | 0.1636 | 14 | 21 | **** |
| South Pacific | 0.0242 | 0.1012 | 25 | 12 | *** |
| Southern Ocean | 0.0112 | 0.1205 | 3 | 1 | N/A |
| Mediterranean | 0.0159 | N/A | 12 | 0 | N/A |
| Red Sea | 0.0088 | N/A | 6 | 0 | N/A |

**Table S3**. Linear regression models to predict the log-transformed transposase abundance in metagenomic samples.

| **Model** | **p for F-test** | **Cumulative R2** |
|---|---|---|
| depth | $< 10^{-10}$ | 0.64 |
| depth + temperature (C°) | 0.48 | 0.65 |
| depth + dissolved oxygen (μmol/Kg) | 0.09 | 0.67 |
| depth + secretory peptidase (%) | $3 \times 10^{-7}$ | 0.70 |
| depth + secretory CAZyme (%) | $4 \times 10^{-9}$ | 0.72 |

**Supplementary Figures:**

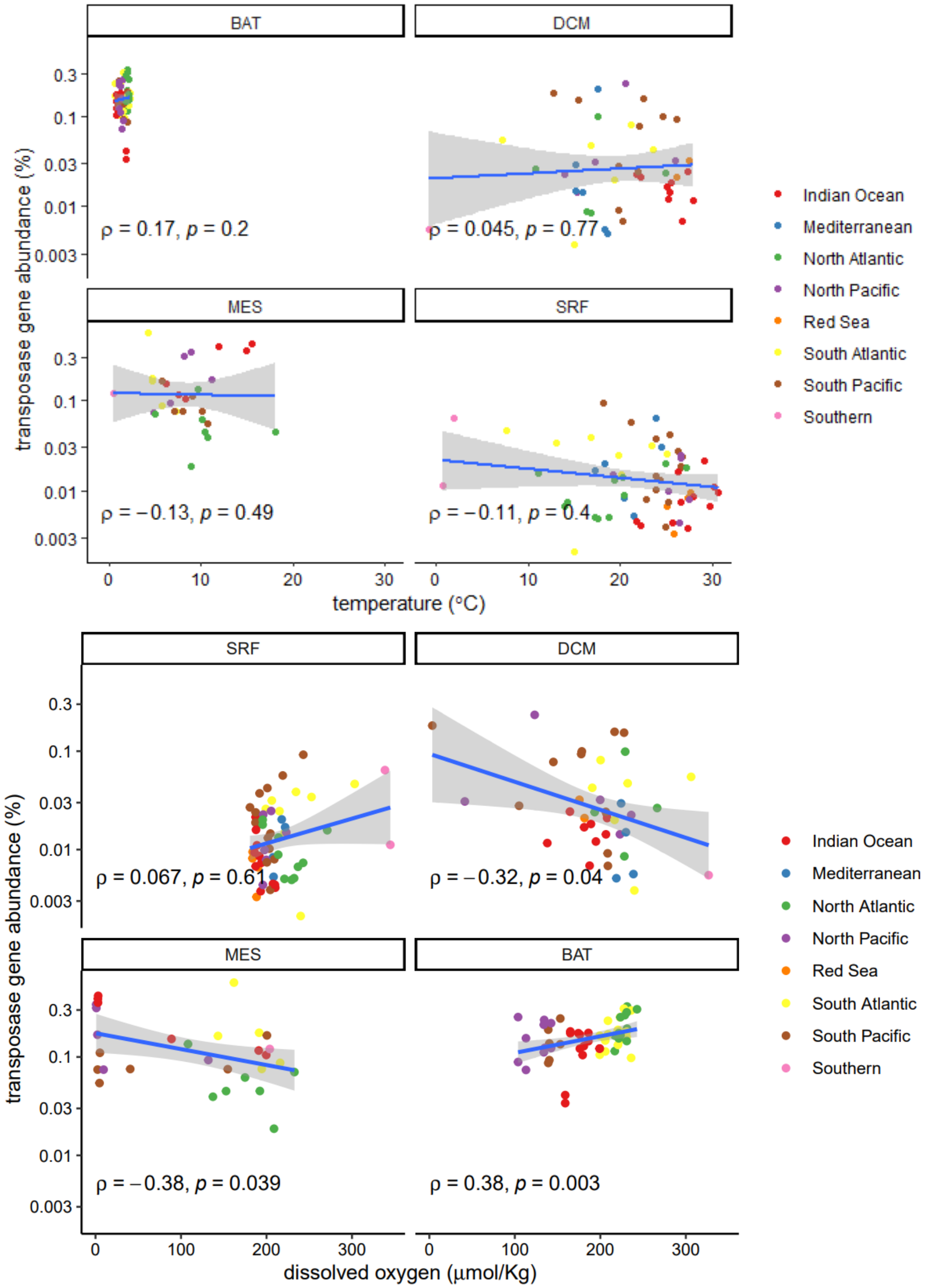


**Fig S1**. **A** The correlation between transposase abundance and temperature in metagenomic samples. **B** The correlation between transposase abundance and the concentration of dissolved oxygen. Each dot is a metagenomic sample. Spearman's correlation coefficient and *p*-value were shown on bottom left.

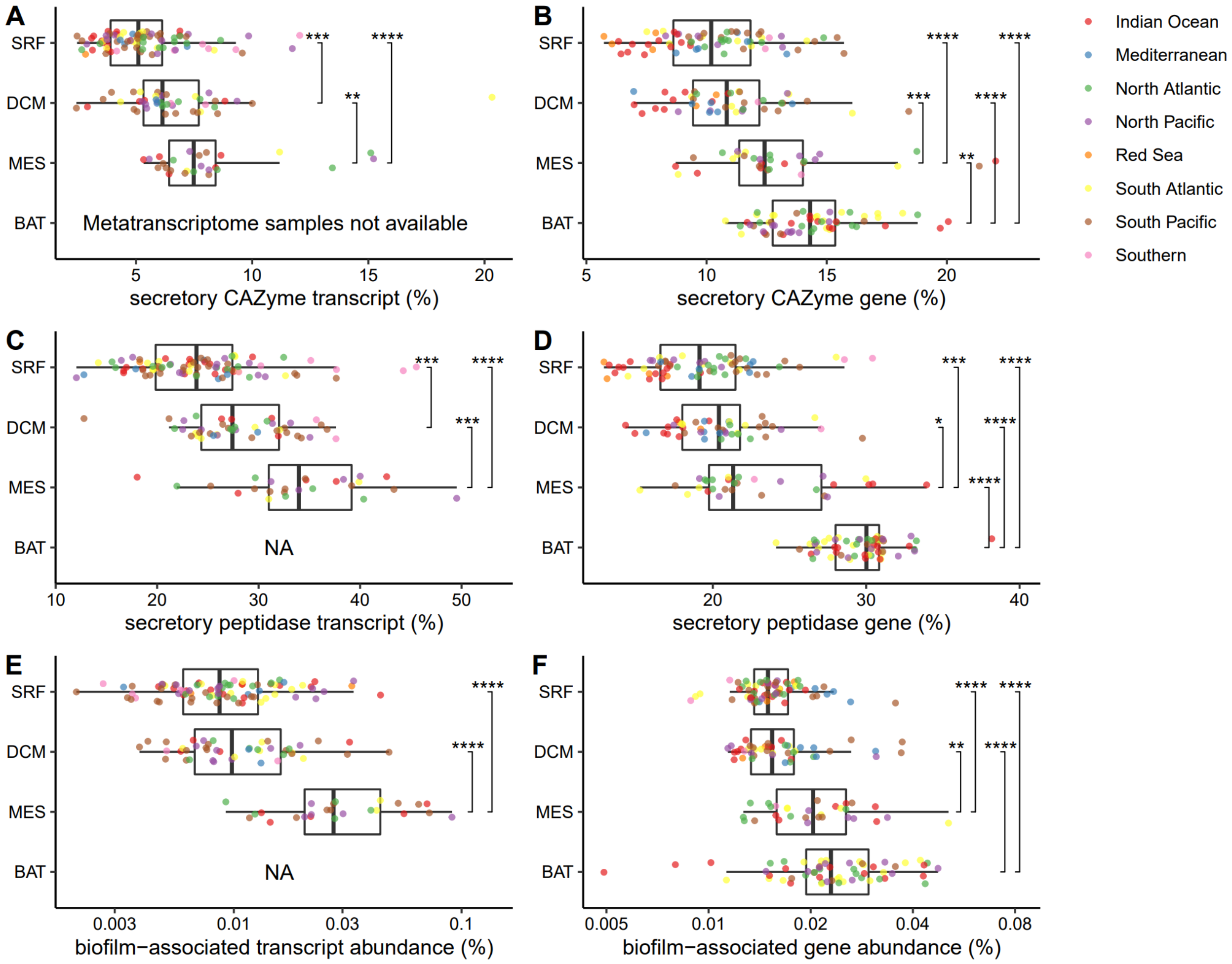


**Fig S2**. Evidence of increasing particle association in microbial communities toward the deep oceans. Each dot is a metatranscriptomic or metagenomic sample, grouped by depth. **A, B** The percentage of secretory CAZyme transcripts/reads in all CAZyme transcripts/reads in each sample. **C, D** The percentage of secretory peptidase transcripts/reads in all peptidase transcripts/reads in each sample. **E, F** The gene and transcript abundance of biofilm-associated ORFs in each sample, separated by depth. *** $p < 0.001$, **** $p < 0.0001$.

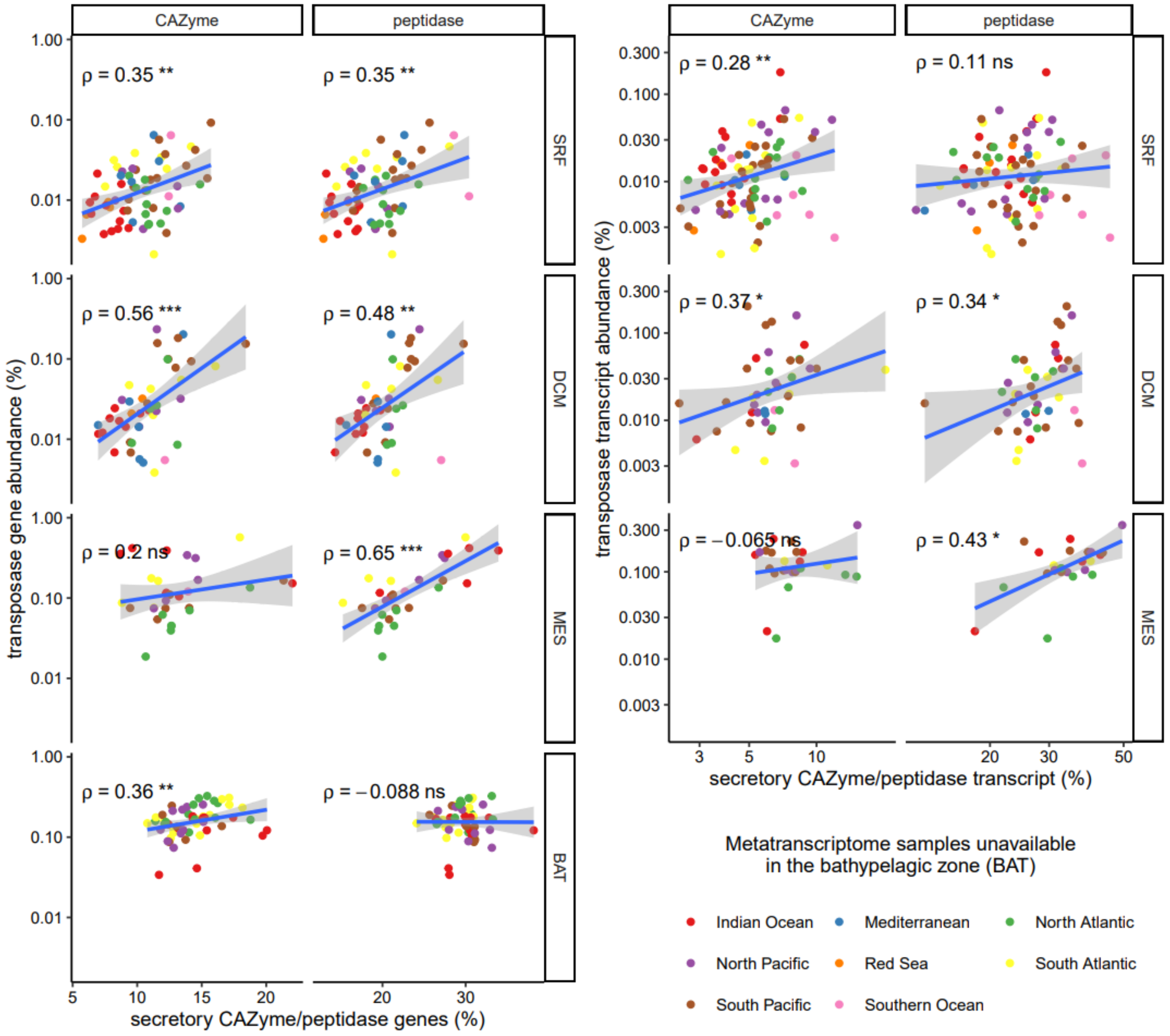


**Fig S3**. The correlation between transposase abundance and the percentage of secretory CAZymes and peptidases ORFs among all secretory CAZymes and peptidases. The Spearman's correlation coefficients, ρ, were shown on the top left. * $p$ < 0.05, ** $p$ < 0.01, *** $p$ < 0.001, **** $p$ < 0.0001.

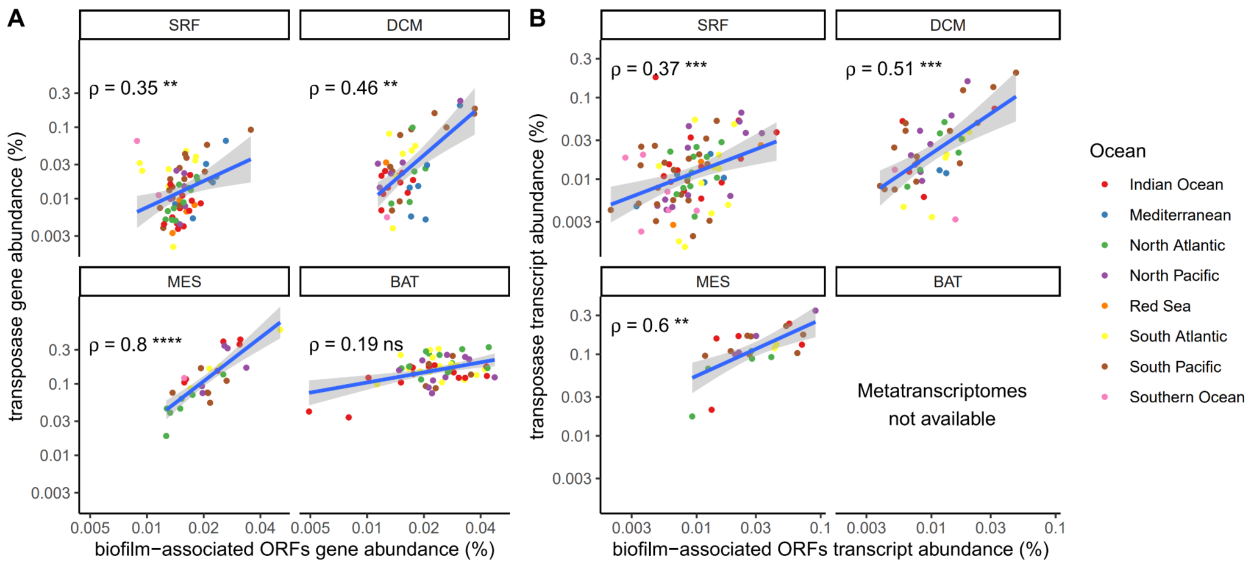


**Fig S4.** The gene potential (DNA) and transcript abundance (RNA) of transposases correlates with those of biofilm genes in all depths except the bathypelagic zone. **A** The correlation between the abundance of transposases and biofilm-associated ORFs in metagenomes (n = 196), separated by depths. **B** The same correlation in metatranscriptomes (n = 181).

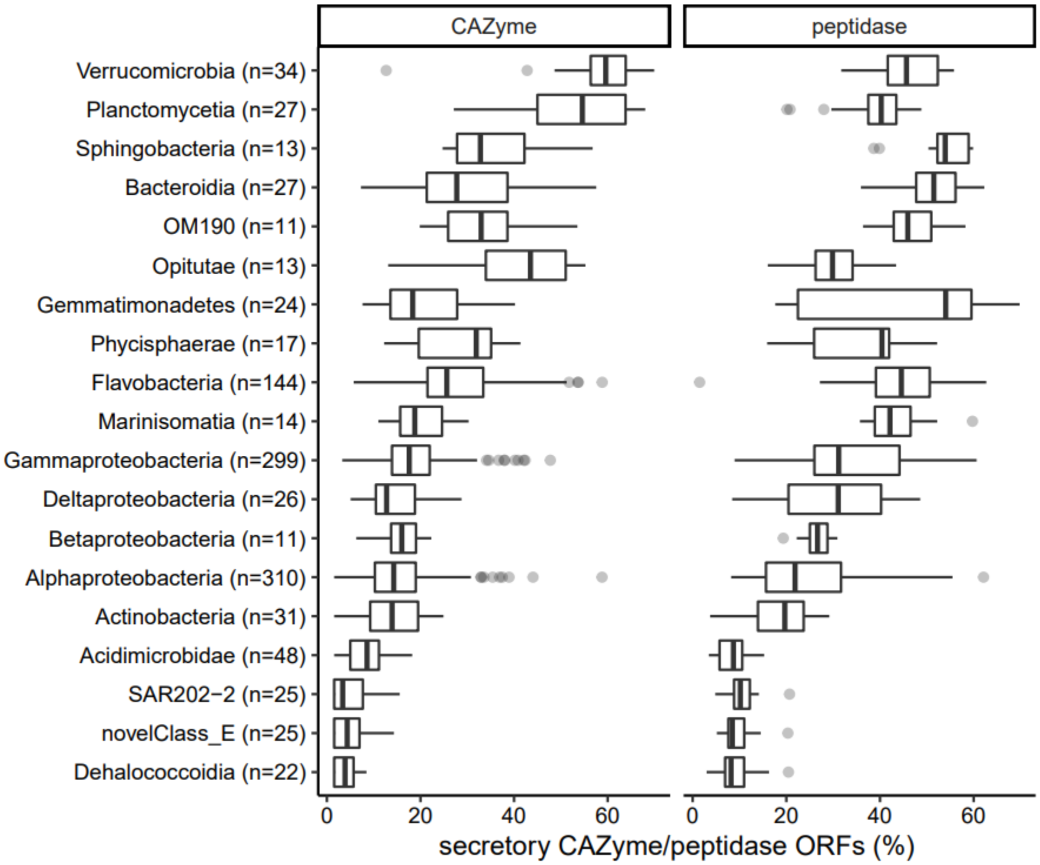


**Fig S5**. Percentages of secretory CAZyme and peptidase ORFs among all CAZyme and peptidase ORFs in MAGs of different taxonomic classes. The percentage is a ratio of ORF counts. Taxa are sorted by the sum of median percentage secretory CAZyme and median percentage of secretory peptidase in a class. The 3 taxonomic classes with lowest percentages of secretory CAZymes and peptidases – *Acidimicrobidae, Delahococcoidia*, and *SAR202-2* – are known to be planktonic (1–4) (there was no pre-existing literature characterizing *novelClass_E*). In contrast, the 3 classes with highest percentages of secretory CAZymes and peptidases – *Verrucomicrobia*, *Planctomycetia*, and *Sphingobacteria* – are known to be particle-associated (4–8).

The extreme ends of the percentage of secretory CAZyme and peptidase reliably predict the lifestyle (planktonic vs. particle-associated) of a given taxon, showing that the percentage of secretory CAZyme and peptidase ORFs is a valuable estimator for the lifestyle of a population.

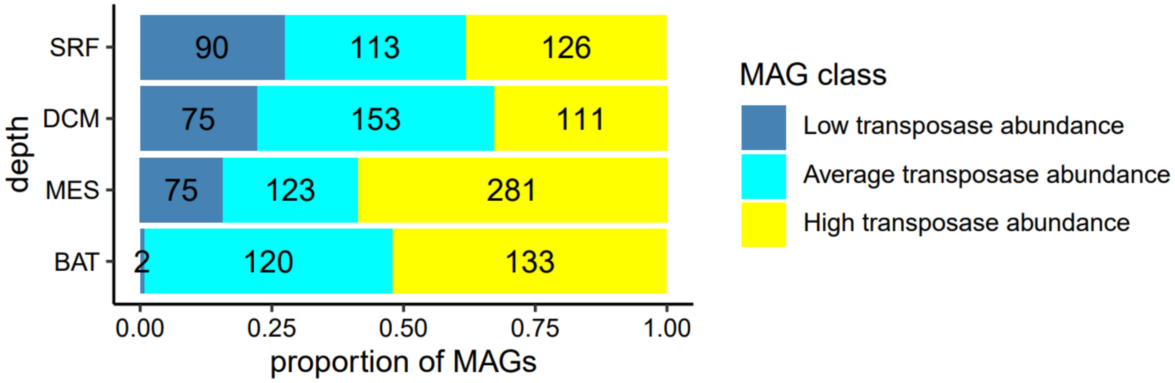


**Fig S6**. Metagenome assembled genomes (MAGs) belonging to a high transposase abundance taxonomic class were more prevalent in the deep ocean. MAGs with no taxonomic prediction were excluded. Classes with **high transposase abundance**: *Alpha-, Beta-, Gammaproteobacteria*, and *Actinobacteria*. Classes with **low transposase abundance**: *Flavobacteria, Acidimicrobidae, SAR202-2*, and *novelClass_E*. All other taxon classes had average transposase abundance.

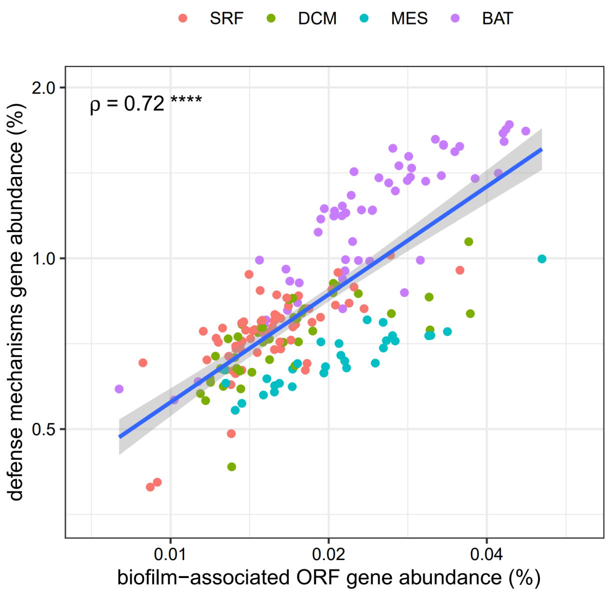


**Fig S7.** The correlation between gene abundances of biofilm-associated ORFs and defense mechanism ORFs in metagenomes, colored by depths. The Spearman's correlation coefficients, $\rho$, are shown on top left. **** $p < 0.0001$.

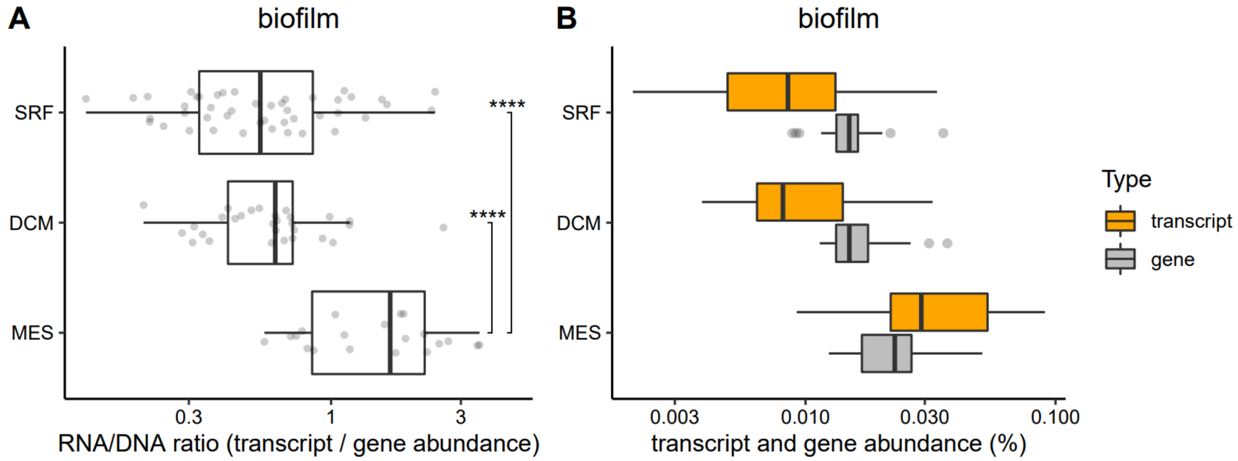


**Fig S8.** Biofilm-associated ORFs were more expressed in deeper waters. **A** Log-transformed RNA/DNA ratio of biofilm-associated ORFs in each sample, separated by depth. The metagenomic and metatranscriptomic abundance of biofilm-associated ORFs were also shown. **** $p < 0.0001$.

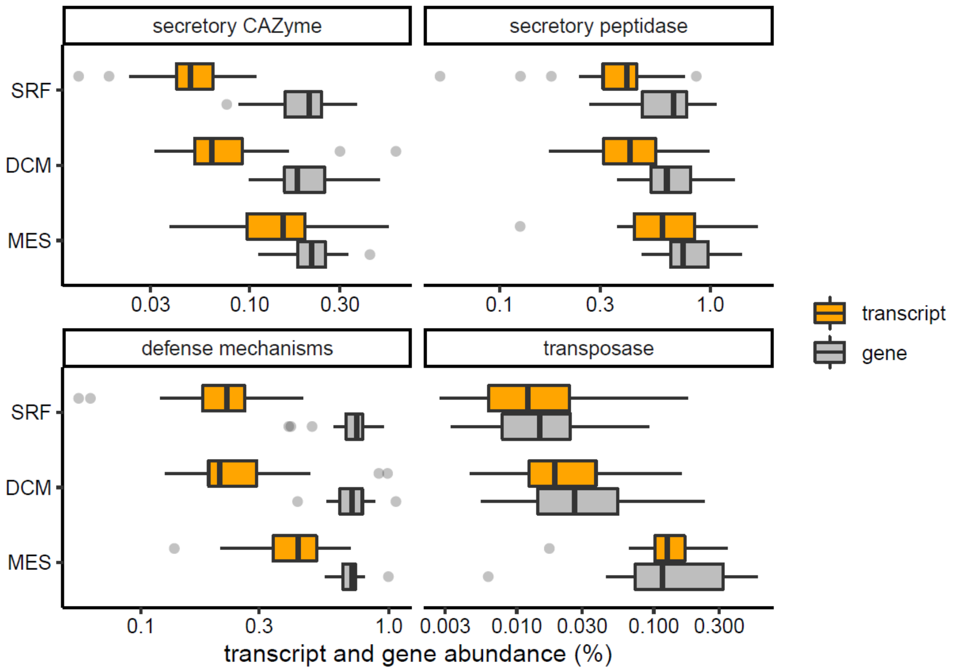


**Fig S9**. The metagenomic and metatranscriptomic abundance of target genes in each depth. Each sample had both metagenome and metatranscriptome sequenced. Target genes were secretory CAZymes, secretory peptidases, genes of the defense mechanisms functional category, and transposases.

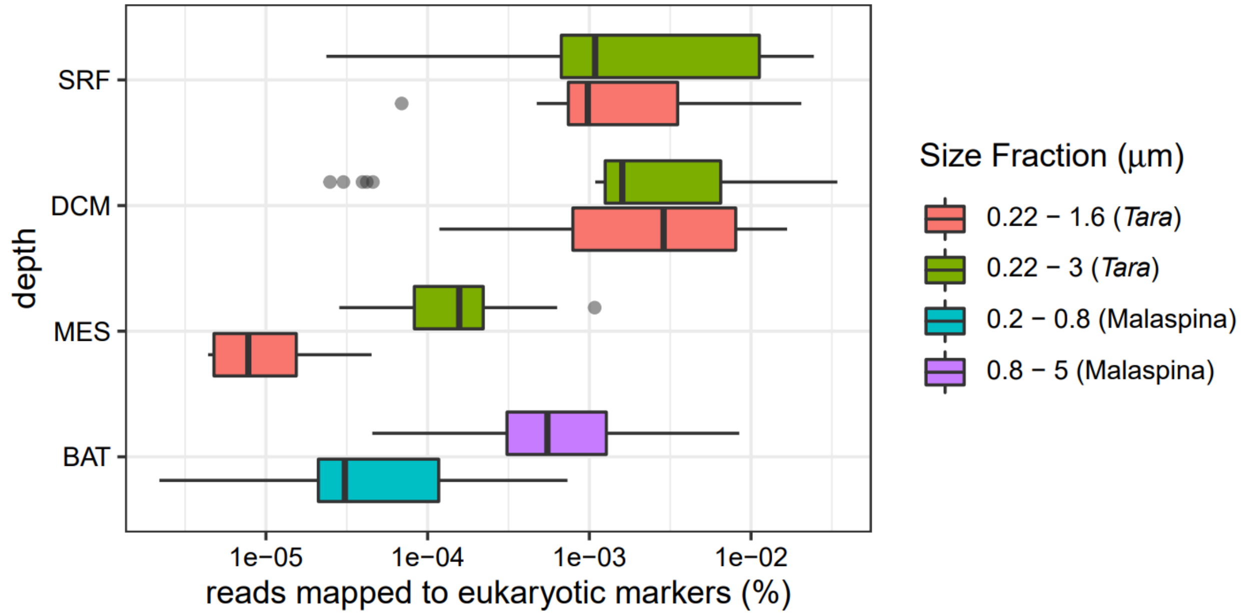


**Fig. S10**. Abundance of reads in metagenomic samples that mapped to eukaryotic marker genes.

#### Supplementary Text

**Setting up anvi'o profiles, obtaining ORF annotations.** We used the command "anvi-profile" from anvi'o pipeline (9) to profile the *Tara* Oceans and Malaspina metagenomes (10, 11). The flag "--profile SCVs" was set to detect single nucleotide variants (SNVs) and single codon variants (SCVs), which were used later to calculate *pN/pS* ratios. During the profiling step, anvi'o identified ORFs from contigs with Prodigal V2.6.3 (9, 12). Functional categories of ORFs were determined with "anvi-run-ncbi-cogs", which used DIAMOND (v0.9.24) (13) and the COG20 database (14, 15).

**Contamination from eukaryotic reads.** Metagenomic samples used here were collected with different filter sizes: samples from the Malaspina expedition were collected on 0.2 – 0.8 µm and 0.8 – 5 µm size filters (11), whereas Tara Oceans samples were collected on 0.22 – 1.6 µm and 0.22 – 3.0 µm size filters (16). Larger size filters could potentially capture more eukaryotic cells, confounding our analysis. Thus, the percentage of eukaryotic reads in our samples was accessed with the software EukDetect (17). The software EukDetect mapped reads to a database of eukaryotic marker genes. We found that samples from the shallow oceans (surface and DCM) showed no difference in the abundance of eukaryotic marker genes between the 0.22 – 1.6 µm and 0.22 – 3.0 µm size filters (**Fig. S10**). Although the abundance of eukaryotic marker genes were different between filter sizes in samples from the deep oceans (mesopelagic and bathypelagic zones), such abundance was small (< 0.01%) and less than those of the shallow samples (**Fig. S10**). Thus, we decided to include all these samples in our analysis.

#### References


1. Mizuno CM, Rodriguez-Valera F, Ghai R. 2015. Genomes of planktonic Acidimicrobiales: widening horizons for marine Actinobacteria by metagenomics. MBio 6.

2. Rowe AR, Lazar BJ, Morris RM, Richardson RE. 2008. Characterization of the community structure of a dechlorinating mixed culture and comparisons of gene expression in planktonic and biofloc-associated "Dehalococcoides" and Methanospirillum species. Appl Environ Microbiol 74:6709–6719.

3. Morris RM, Rappé MS, Urbach E, Connon SA, Giovannoni SJ. 2004. Prevalence of the Chloroflexi-related SAR202 bacterioplankton cluster throughout the mesopelagic zone and deep ocean. Appl Environ Microbiol 70:2836–2842.

4. Leu AO, Eppley JM, Burger A, DeLong EF. 2022. Diverse Genomic Traits Differentiate Sinking-Particle-Associated versus Free-Living Microbes throughout the Oligotrophic Open Ocean Water Column. MBio 13:e0156922.

5. Bengtsson MM, Øvreås L. 2010. Planctomycetes dominate biofilms on surfaces of the kelp Laminaria hyperborea. BMC Microbiol 10:261.

6. Lage OM, Bondoso J. 2014. Planctomycetes and macroalgae, a striking association. Front Microbiol 5:267.

7. Cardman Z, Arnosti C, Durbin A, Ziervogel K, Cox C, Steen AD, Teske A. 2014. Verrucomicrobia are candidates for polysaccharide-degrading bacterioplankton in an arctic fjord of Svalbard. Appl Environ Microbiol 80:3749–3756.

8. Ganesh S, Parris DJ, DeLong EF, Stewart FJ. 2014. Metagenomic analysis of size-fractionated picoplankton in a marine oxygen minimum zone. ISME J 8:187–211.

9. Eren AM, Kiefl E, Shaiber A, Veseli I, Miller SE, Schechter MS, Fink I, Pan JN, Yousef M, Fogarty EC, Trigodet F, Watson AR, Esen ÖC, Moore RM, Clayssen Q, Lee MD, Kivenson V, Graham ED, Merrill BD, Karkman A, Blankenberg D, Eppley JM, Sjödin A, Scott JJ, Vázquez-Campos X, McKay LJ, McDaniel EA, Stevens SLR, Anderson RE, Fuessel J, Fernandez-Guerra A, Maignien L, Delmont TO, Willis AD. 2021. Community-led, integrated, reproducible multi-omics with anvi'o. Nat Microbiol 6:3–6.

10. Tully BJ, Graham ED, Heidelberg JF. 2018. The reconstruction of 2,631 draft metagenome-assembled genomes from the global oceans. Scientific Data 5:1–8.

11. Acinas SG, Sánchez P, Salazar G, Cornejo-Castillo FM, Sebastián M, Logares R, Royo-Llonch M, Paoli L, Sunagawa S, Hingamp P, Ogata H, Lima-Mendez G, Roux S, González JM, Arrieta JM, Alam IS, Kamau A, Bowler C, Raes J, Pesant S, Bork P, Agustí S, Gojobori T, Vaqué D, Sullivan MB, Pedrós-Alió C, Massana R, Duarte CM, Gasol JM. 2021. Deep ocean metagenomes provide insight into the metabolic architecture of bathypelagic microbial communities. Communications Biology 4:1–15.

12. Hyatt D, Chen G-L, Locascio PF, Land ML, Larimer FW, Hauser LJ. 2010. Prodigal: prokaryotic gene recognition and translation initiation site identification. BMC Bioinformatics 11:119.

13. Buchfink B, Reuter K, Drost H-G. 2021. Sensitive protein alignments at tree-of-life scale using DIAMOND. Nat Methods 18:366–368.

14. Galperin MY, Wolf YI, Makarova KS, Vera Alvarez R, Landsman D, Koonin EV. 2020. COG database update: focus on microbial diversity, model organisms, and widespread pathogens. Nucleic Acids Res 49:D274–D281.

15. Tatusov RL. 2000. The COG database: a tool for genome-scale analysis of protein

functions and evolution. Nucleic Acids Research https://doi.org/10.1093/nar/28.1.33.

16. Sunagawa S, Acinas SG, Bork P, Bowler C, Tara Oceans Coordinators, Eveillard D, Gorsky G, Guidi L, Iudicone D, Karsenti E, Lombard F, Ogata H, Pesant S, Sullivan MB, Wincker P, de Vargas C. 2020. Tara Oceans: towards global ocean ecosystems biology. Nat Rev Microbiol 18:428–445.

17. Lind AL, Pollard KS. 2021. Accurate and sensitive detection of microbial eukaryotes from whole metagenome shotgun sequencing. Microbiome.
